# Multiple C2 domains and Transmembrane region Proteins (MCTPs) tether membranes at plasmodesmata

**DOI:** 10.1101/423905

**Authors:** Marie L. Brault, Jules D. Petit, Françoise Immel, William J. Nicolas, Lysiane Brocard, Amélia Gaston, Mathieu Fouché, Timothy J. Hawkins, Jean-Marc Crowet, Magali S. Grison, Max Kraner, Vikram Alva, Stéphane Claverol, Magali Deleu, Laurence Lins, Jens Tilsner, Emmanuelle M. Bayer

## Abstract

In eukaryotes, membrane contact sites (MCS) allow direct communication between organelles. Plants have evolved unique MCS, the plasmodesmata intercellular pores, which combine endoplasmic reticulum (ER) - plasma membrane (PM) contacts with regulation of cell-to-cell signalling. The molecular mechanism and function of membrane tethering within plasmodesmata remains unknown.

Here we show that the Multiple C2 domains and Transmembrane region Protein (MCTP) family, key regulators of cell-to-cell signalling in plants, act as ER - PM tethers specifically at plasmodesmata. We report that MCTPs are core plasmodesmata proteins that insert into the ER via their transmembrane region whilst their C2 domains dock to the PM through interaction with anionic phospholipids. A *mctp3/4* loss-of-function mutant induces plant developmental defects while MCTP4 expression in a yeast Δtether mutant partially restores ER-PM tethering. Our data suggest that MCTPs are unique membrane tethers controlling both ER-PM contacts and cell-cell signalling.

## Introduction

Intercellular communication is essential for the establishment of multicellularity, and evolution gave rise to distinct mechanisms to facilitate this process. Plants have developed singular cell junctions -the plasmodesmata-which span the cell wall and interconnect nearly every single cell, establishing direct membrane and cytoplasmic continuity throughout the plant body (Tilsner *et al*, 2016). Plasmodesmata are indispensable for plant life. They control the flux of non-cell-autonomous signals such as transcription factors, small RNAs, hormones and metabolites during key growth and developmental events (Gallagher *et al*, 2014; Tilsner *et al*, 2016; Vatén *et al*, 2011; Carlsbecker *et al*, 2010; Benitez-Alfonso *et al*, 2013; Wu *et al*, 2016; Han *et al*, 2014; Daum *et al*, 2014; Nakajima *et al*, 2001; Xu *et al*, 2011; Ross-elliott *et al*, 2017). Over the past few years, plasmodesmata have emerged as key components of plant defence signalling (Faulkner *et al*, 2013; Wang *et al*, 2013; Lim *et al*, 2016). Mis-regulation of plasmodesmata function can lead to severe defects in organ growth and tissue patterning but also generate inappropriate responses to biotic and abiotic stresses (Wu *et al*, 2016; Wong *et al*, 2016; Han *et al*, 2014; Sager & Lee, 2014; Caillaud *et al*, 2014; Faulkner *et al*, 2013). Plasmodesmata not only serve as conduits, but act as specialised signalling hubs, capable of generating and/or relaying signals from cell-to-cell through plasmodesmata-associated receptor activity (Stahl & Faulkner, 2016; Stahl *et al*, 2013; Vaddepalli *et al*, 2014; Lee, 2015).

Plasmodesmata are structurally unique (Nicolas *et al*, 2017b; Tilsner *et al*, 2011). They contain a strand of ER, continuous through the pores, tethered extremely tightly (~10 nm) to the PM by spoke-like elements (Ding *et al*, 1992; Nicolas *et al*, 2017a) whose function and identity is unknown. Inside plasmodesmata, specialised subdomains of the ER and the PM coexist, each being characterised by a unique set of lipids and proteins, both critical for proper function (Bayer *et al*, 2014; Grison *et al*, 2015; Thomas *et al*, 2008a; Simpson *et al*, 2009; Zavaliev *et al*, 2016; Knox *et al*, 2015; Faulkner *et al*, 2013; Benitez-Alfonso *et al*, 2013). Where it enters the pores, the ER becomes constricted to a 15 nm tube (the desmotubule) leaving little room for lumenal trafficking. According to current models, transfer of molecules occurs in the cytoplasmic sleeve between the ER and the PM. Constriction of this gap, by the deposition of callose, is assumed to be the main regulator of the pore size exclusion limit (Vatén *et al*, 2011; Zavaliev *et al*, 2011). Recent work however, suggests a more complex picture where plasmodesmal ER-PM gap is not directly related to the pore permeability and may play additional roles (Nicolas *et al*, 2017a, 2017b). Newly formed plasmodesmata (type I) exhibit such close contact (~2-3nm) between the PM and the ER, that no electron-lucent cytoplasmic sleeve is observed (Nicolas *et al*, 2017a). During subsequent cell growth and differentiation the pore widens, separating the two membranes, which remain connected by visible electron-dense spokes, leaving a cytosolic gap (type II). This transition has been proposed to be controlled by protein-tethers acting at the ER-PM interface (Bayer *et al*, 2017; Nicolas *et al*, 2017b). Counterintuitively, type I plasmodesmata with no apparent cytoplasmic sleeve are open to macromolecular trafficking and recent data indicate that tight ER-PM contacts may in fact favour transfer of molecules from cell-to-cell (Nicolas *et al*, 2017a).

The close proximity of the PM and ER within the pores, and the presence of tethers qualifies plasmodesmata as a specialised type of ER-to-PM membrane contact site (MCS) (Tilsner *et al*, 2016; Bayer *et al*, 2017). MCS are structures found in all eukaryotic cells which function in direct inter-organellar signalling by promoting fast, non-vesicular transfer of molecules and allowing collaborative action between the two membranes (Burgoyne *et al*, 2015; Prinz, 2014; Phillips & Voeltz, 2016; Gallo *et al*, 2016; Eden *et al*, 2010, 2016; Ho *et al*, 2016; Chang *et al*, 2013; Kim *et al*, 2015; Petkovic *et al*, 2014; Zhang *et al*, 2005; Omnus *et al*, 2016). In yeast and mammalian, MCS protein tethers are known to physically bridge the two organelles, to control the intermembrane gap and participate in organelle cross-talk. Their molecular identity/specificity dictate structural and functional singularity to different types of MCS (Eisenberg-Bord *et al*, 2016; Henne *et al*, 2015). To date, the plasmodesmal membrane tethers remain unidentified, but by analogy to other types of MCS it seems likely that they play important roles in plasmodesmal structure and function, and given their unique position within a cell-to-cell junction may link intra- and intercellular communication.

Here, we have reduced the complexity of the previously published *Arabidopsis* plasmodesmal proteome (Fernandez-Calvino *et al*, 2011) through the combination of a refined purification protocol (Faulkner & Bayer, 2017) and semi-quantitative proteomics, to identify ~120 proteins highly enriched in plasmodesmata, and identify tether candidates. Amongst the most abundant plasmodesmal proteins, members of the Multiple C2 domains and Transmembrane region Proteins (MCTPs) were enriched in post-cytokinetic plasmodesmata with tight ER-PM gap compared to mature plasmodesmata with wider gap and sparse spokes, and exhibit the domain architecture characteristic of membrane tethers, with multiple lipid-binding C2 domains in the N-terminal, and multiple transmembrane domains in the C-terminal region. Using live cell imaging, molecular dynamics, and yeast complementation, we show that MCTP properties are consistent with a role in ER-PM membrane tethering at plasmodesmata. As several MCTP members have been identified as important components of plant intercellular signalling (Liu *et al*, 2012, 2018; Vaddepalli *et al*, 2014), our data suggest a link between inter-organelle contacts at plasmodesmata and inter-cellular communication in plants.

## RESULTS

### Identification of plasmodesmal ER-PM tethering candidates

To identify putative plasmodesmal MCS tethers, we decided to screen the plasmodesmata proteome for ER-associated proteins (a general trait of ER-PM tethers (Henne *et al*, 2015; Eisenberg-Bord *et al*, 2016)) with structural features enabling bridging across two membranes. Published plasmodesmata proteome reported the identification of more than 1400 proteins in *Arabidopsis* (Fernandez-Calvino *et al*, 2011), making the discrimination of true plasmodesmata-associated from contaminant proteins a major challenge. To reduce the proteome complexity and identify core plasmodesmata proteins, we used a refined plasmodesmata purification technique (Faulkner & Bayer, 2017) together with label-free comparative quantification (Supplementary Fig. 1a). Plasmodesmata and likely contaminant fractions, namely the PM, microsomal, total cell and cell wall fractions were purified from six-day old *Arabidopsis* suspension culture cells and simultaneously analysed by liquid-chromatography tandem mass-spectrometry (LC-MS/MS). For each protein identified, its relative enrichment in the plasmodesmata fraction versus “contaminant” fractions was determined (Supplementary Fig. 1b; Supplementary Table 1). Enrichment ratios for selecting plasmodesmal-candidates was set based on previously characterised plasmodesmal proteins (see M&M for details). This refined proteome dataset was reduced to 115 unique proteins, cross-referenced with two published ER-proteomes (Nikolovski *et al*, 2012; Dunkley *et al*, 2006) and used as a basis for selecting MCS-relevant candidates.

Alongside, we also analysed changes in protein abundance during the ER-PM tethering transition from very tight contacts in post-cytokinetic plasmodesmata (type I) to larger ER-PM gap and sparse tethers in mature plasmodesmata (type II) (Nicolas *et al*, 2017a). For this we obtained a similar semi-quantitative proteome from four and seven-day old culture cells, enabling a comparison of plasmodesmata composition during the tethering transition (Nicolas *et al*, 2017a) (Supplementary Fig. 2).

A survey of our refined proteome identified several members of the Multiple C2 domains and Transmembrane region Proteins (MCTPs) family, namely AtMCTP3-7, 9, 10, 14-16, as both abundant and highly enriched at plasmodesmata (Supplementary Fig. 1b, Supplementary Table 1). In addition to being “core” plasmodesmata-associated proteins, our data also suggests that MCTPs are differentially regulated during the ER-PM tethering transition from post-cytokinetic to mature plasmodesmata (Nicolas *et al*, 2017a) (Supplementary Fig. 2). Amongst the 47 plasmodesmal proteins differentially enriched, all MCTPs were more abundant (1.4 to 3.6 times) in type I (tight ER-PM contacts) compared with type II (open cytoplasmic sleeves) plasmodesmata (Supplementary Fig. 2).

### MCTPs are ER-associated proteins that stably cluster at plasmodesmata and present structural features of membrane tethers

MCTPs are structurally reminiscent of the ER-PM tether families of mammalian extended-Synaptotagmins (HsE-Syts) and plant *Arabidopsis* Synaptotagmins (AtSYTs) (Pérez-Sancho *et al*, 2015b; Giordano *et al*, 2013), possessing lipid-binding C2 domains at one end and multiple transmembrane domains (TMDs) at the other, a domain organization consistent with the function of membrane tethers (Supplementary Fig. 3). Unlike HsE-Syts and AtSYTs, the transmembrane region of MCTPs is located at the C-terminus and three to four C2 domains at the N-terminus (Fig. 1a; Supplementary Fig. 3). Two members of the *Arabidopsis* MCTP family, AtMCTP1/Flower locus T Interacting Protein (FTIP) and AtMCTP15/QUIRKY (QKY) have previously been localised to plasmodesmata in *Arabidopsis* and implicated in cell-to-cell trafficking of developmental signals (Vaddepalli *et al*, 2014; Liu *et al*, 2012). However, two recent studies indicate that other MCTP members, including AtMCTP3, 4, 9, which show high plasmodesmata-enrichment in our proteome, do not associate with the pores *in vivo* (Liu *et al*, 2017, 2018).

**Figure 1.**
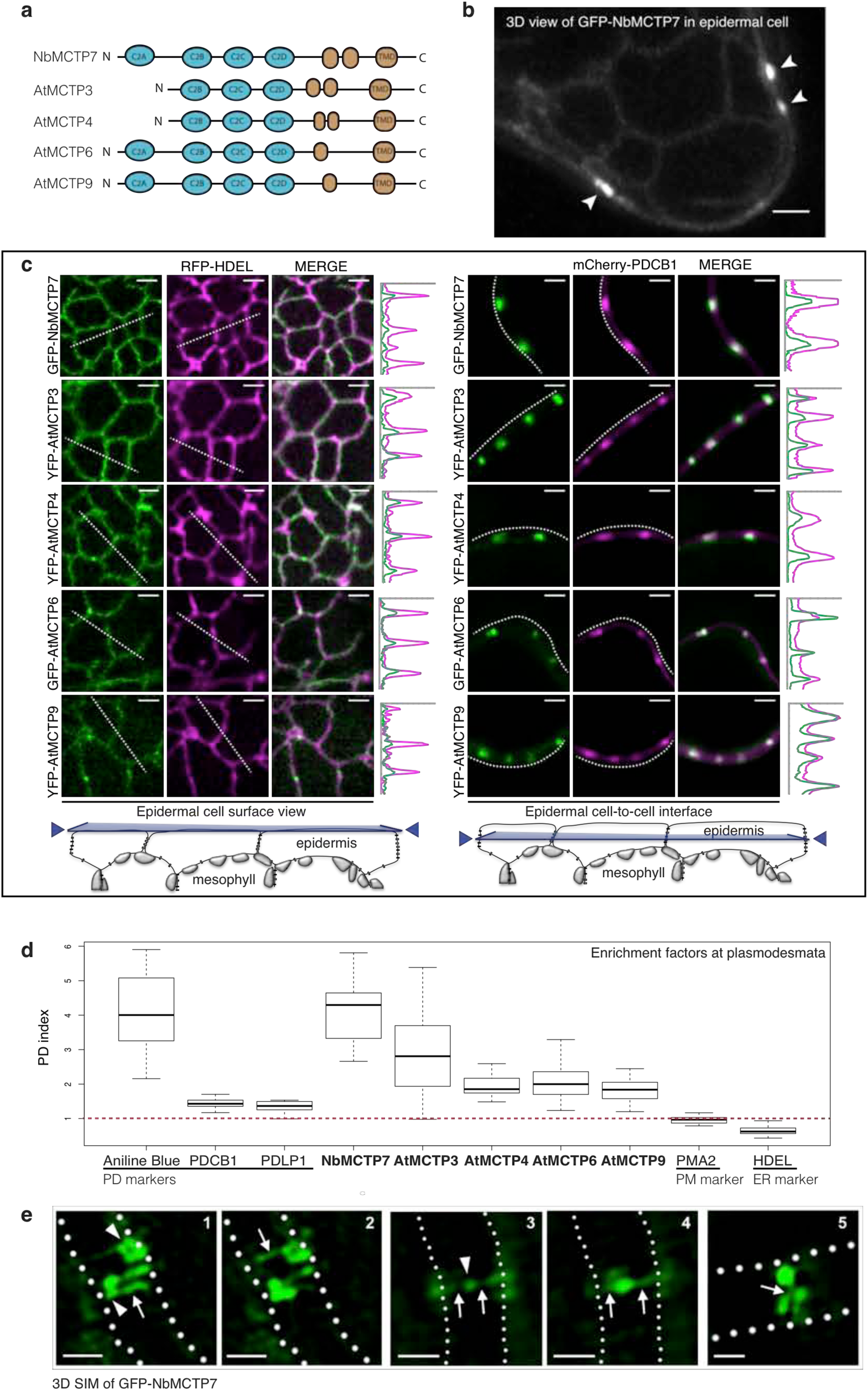
MCTPs are ER-associated proteins that cluster at plasmodesmata. Localisation of AtMCTP3, 4, 6, 9 and NbMCTP7 in *N. benthamiana* epidermal cells visualised by confocal microscopy. MCTPs were tagged at their N-terminus with YFP or GFP and expressed transiently under 35S (NbMCTP7) or UBIQUITIN10 promoters (AtMCTP3, 4, 6 and 9). **a**, Schematic representation of MCTP domain organisation, with three to four C2 domains in N-terminal and multiple transmembrane domains (TMD) at the C-terminus. **b**, GFP-NbMCTP7 associates with punctae at the cell periphery (white arrowheads) and labels a reticulated network at the cell surface resembling the cortical ER. Maximum projection of z-stack. Scale bar, 2 μm **c**, Single optical sections at cell surface (left) or cell-to-cell interface (right), showing the co-localisation between MCTPs and the ER-marker RFP-HDEL (left) and the plasmodesmata marker mCherry-PDCBl (right). Intensity plots along the white dashed lines are shown for each co-localisation pattern. Scale bars, 2 μm. **d**, The plasmodesmata (PD) index of individual MCTPs is above 1 (red dashed line), and similar to known plasmodesmata markers (aniline blue, PDCB1, PDLP1) confirming enrichment at plasmodesmata. By comparison the PM-localised proton pump ATPase PMA2 and the ER marker HDEL that are not enriched at plasmodesmata and have a PD-index below 1. **e**, 3D-SIM images (individual z-sections) of GFP-NbMCTP7 within three different pit fields (panels 1-2, 3-4 and 5, respectively) showing fluorescence signal continuity throughout the pores, enrichment at plasmodesmal neck regions (1-2, arrowheads in 1), central cavity (3-4, arrowhead in 3) and branching at central cavity (5, arrow). Dashed lines indicate position of cell wall borders. Scale bars, 500 nm.

We investigated the *in vivo* localisation of MCTPs identified in our proteomic screen by transiently expressing N-terminal fusions fluorescent proteins in *Nicotiana benthamiana* leaves. As the MCTP family is conserved in *N. benthamiana* (Supplementary Fig. 4) and to avoid working in a heterologous system we also examined the localisation of NbMCTP7, whose closest homolog in *Arabidopsis* was also identified as highly-enriched in plasmodesmata fractions (AtMCTP7; Supplementary Fig. 1). Confocal imaging showed that all selected MCTPs, namely AtMCTP3, 4, 6, 9 and NbMCTP7, displayed a similar subcellular localisation, with a faint ER-like network at the cell surface and a punctate distribution along the cell periphery at sites of epidermal cell-to-cell junctions (Fig. 1b, c). Time-lapse imaging showed that peripheral fluorescent punctae were immobile, which contrasted with the high mobility of the ER-like network (Supplementary Mov. 1). Colocalisation with RFP-HDEL confirmed MCTPs association with the cortical ER, while the immobile spots at the cell periphery perfectly co-localised with plasmodesmal markers (mCherry-PDCB1; (Simpson *et al*, 2009; Grison *et al*, 2015); Fig. 1c). Co-labelling with general ER-PM tethers such as VAP27.1-RFP and SYT1-RFP (Pérez-Sancho *et al*, 2015a; Wang *et al*, 2014), showed partial overlap with GFP-NbMCTP7, while co-localisation with mCherry-PDCBl was significantly higher (Supplementary Fig. 5). To further quantify and ascertain MCTP association with plasmodesmata, we measured a plasmodesmal enrichment ratio, hereafter named “plasmodesmata-index”. For this we calculated fluorescence intensity at plasmodesmata pit-fields (indicated by mCherry-PDCB1 or aniline blue) *versus* cell periphery. All MCTPs tested displayed a high plasmodesmata-index, ranging from 1.85 to 4.15, similar to PDLP1 (1.36) and PDCB1 (1.45) two-well established plasmodesmata markers (Thomas *et al*, 2008b; Simpson *et al*, 2009) (Fig. 1d), confirming enrichment of MCTPs at pit-fields. When stably expressed in *Arabidopsis thaliana* under the moderate promoter UBIQUITIN 10 or 35S promoters AtMCTP4, AtMCTP6 and AtMCTP9 were found mainly restricted to plasmodesmata (Supplementary Fig. 6a, white arrows), as indicated by an increase of their plasmodesmata-index compared with transient expression in *N. benthamiana* (Supplementary Fig. 6b). A weak but consistent ER localisation was also visible in stably transformed *Arabidopsis* (Supplementary Fig. 6a red stars).

To get a better understanding of MCTP distribution within the plasmodesmal pores, we further analysed transiently-expressed GFP-NbMCTP7 by 3D structured illumination superresolution microscopy (3D-SIM) (Fitzgibbon *et al*, 2010) (Fig. 1e). We found that NbMCTP7 is associated with all parts of plasmodesmata including the neck regions and central cavity, as well as showing continuous fluorescence throughout the pores. In some cases, lateral branching of plasmodesmata within the central cavity was resolved. The very faint continuous fluorescent threads connecting neck regions and central cavity correspond to the narrowest regions of the pores and may indicate association with the central desmotubule (Fig. 1e, white arrows).

Using Fluorescence Recovery After Photobleaching (FRAP) we then assessed the mobility of NbMCTP7. We found that, when associated with the cortical ER the fluorescence recovery rate of GFP-NbMCTP7 was extremely fast and similar to RFP-HDEL with half-times of 1.16 sec and 0.99 sec, respectively (Fig. 2). By contrast, when GFP-NbMCTP7 was associated with plasmodesmata, the recovery rate slowed down to a half-time of 4.09 sec, indicating restricted mobility, though still slightly faster than for the cell wall-localised plasmodesmal marker mCherry-PDCB1 (5.98 sec). Overall, these results show that NbMCTP7 mobility is high at the cortical ER but becomes restricted inside the pores.

**Figure 2.**
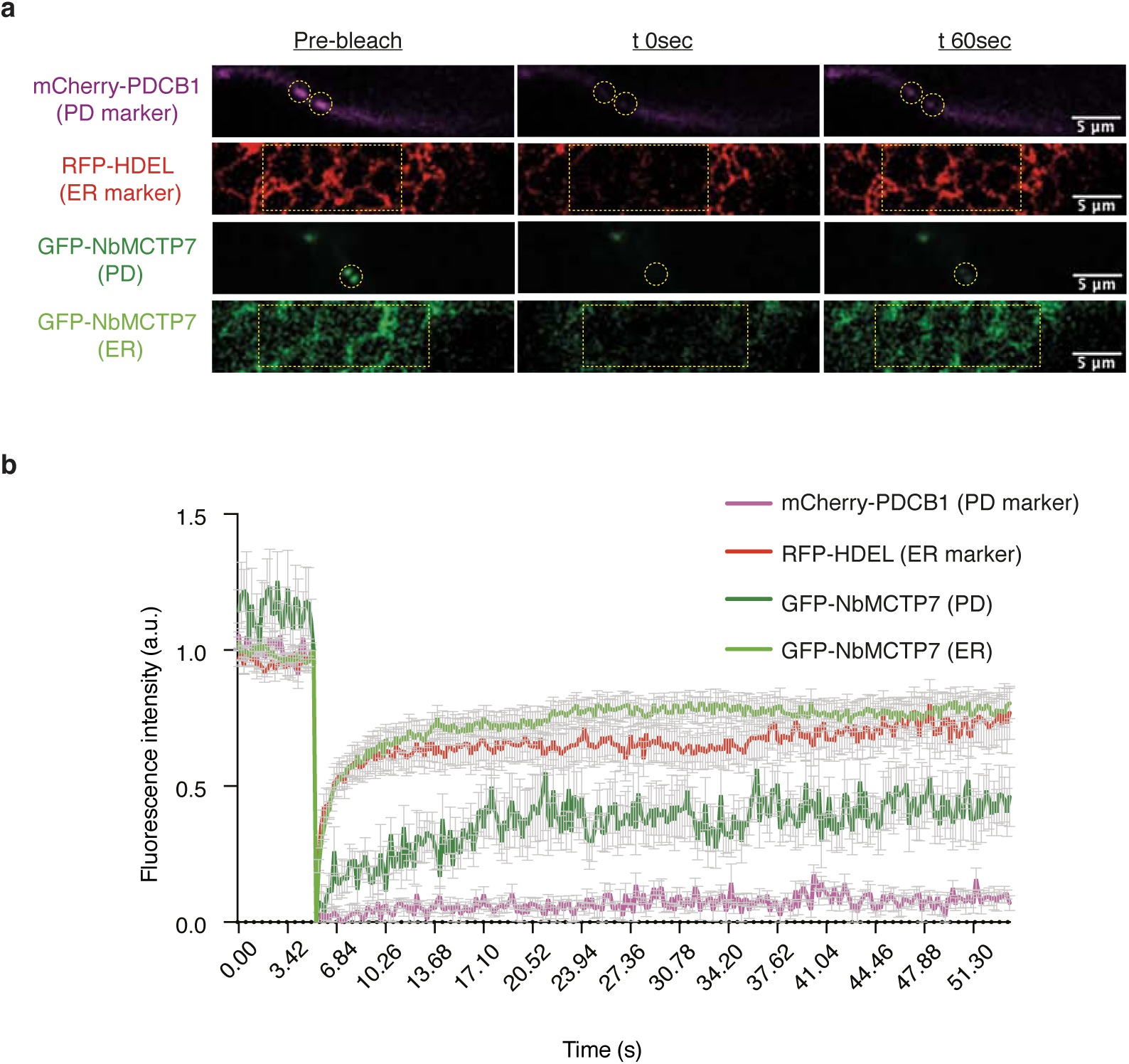
NbMCTP7 mobility at plasmodesmata is reduced compared to cortical ER. FRAP analysis of NbMCTP7 in *N. benthamiana* leaf epidermal cells. **a**, Representative prebleach and postbleach images for mCherry-PDCBl (purple; plasmodesmata marker), RFP-HDEL (red; ER marker) and GFP-NbMCTP7 at plasmodesmata (dark green) and at the cortical ER (light green). Yellow dashed boxes or circles indicate the bleach region. **b**, FRAP comparing the mobility of GFP-NbMCTP7 at plasmodesmata (dark green) and at the cortical ER (light green) to that of RFP-HDEL (red) and mCherry-PDCBl (purple). NbMCTP7 is highly mobile when associated with the ER as indicated by fast fluorescent recovery but shows reduced mobility when associated with plasmodesmata. Data are averages of at least 3 separate experiments, and error bars indicate standard error.

From our data we concluded that MCTPs are ER-associated proteins, whose members specifically and stably cluster at plasmodesmata. They display the structural features required for ER-PM tethering and are differentially associated with the pores during the transition in ER-PM contacts.

### AtMCTP4 is a core plasmodesmata-associated protein and loss-of-function *mctp3/mctp4* double mutants show pleiotropic developmental defects

We next focused on AtMCTP4, which according to our proteomic screen qualifies as a core plasmodesmal constituent considering that it is one of the most abundant proteins in our refined proteome (Supplementary Table 1). The implication of AtMCTP4 association with plasmodesmata is that the protein contributes functionally to cell-to-cell signalling. Given the importance of plasmodesmata in tissue patterning and organ growth, a loss-of-function mutant is expected to show defects in plant development. We first obtained T-DNA insertion lines for AtMCTP4 and its closest homolog AtMCTP3, which share 92.8% identity and 98.7% similarity in amino acids with AtMCTP4, but both single knockouts showed no apparent phenotypic defects (data not shown). We therefore generated an *Atmctp3/Atmctp4* double mutant, which presented pleiotropic developmental defects with a severely dwarfed and bushy phenotype, twisted leaves with increased serration (Fig. 3 a-d), and multiple inflorescences (not shown). Whilst preparing this manuscript another paper describing the *Atmctp3/Atmctp4* mutant was published (Liu *et al*, 2018), reporting similar developmental defects. We noted additional phenotypic defects in particular aberrant pattern in the root apical meristem organisation specifically within the quiescent center (QC) (Fig. 3e). Instead of presenting the typical four-cell layer organisation, we observed aberrant cell division pattern in *Atmctp3/Atmctp4* mutant with asymmetrical division in the QC, suggesting that both proteins may play a general role in cell stem niche maintenance (Liu *et al*, 2018).

**Figure 3.**
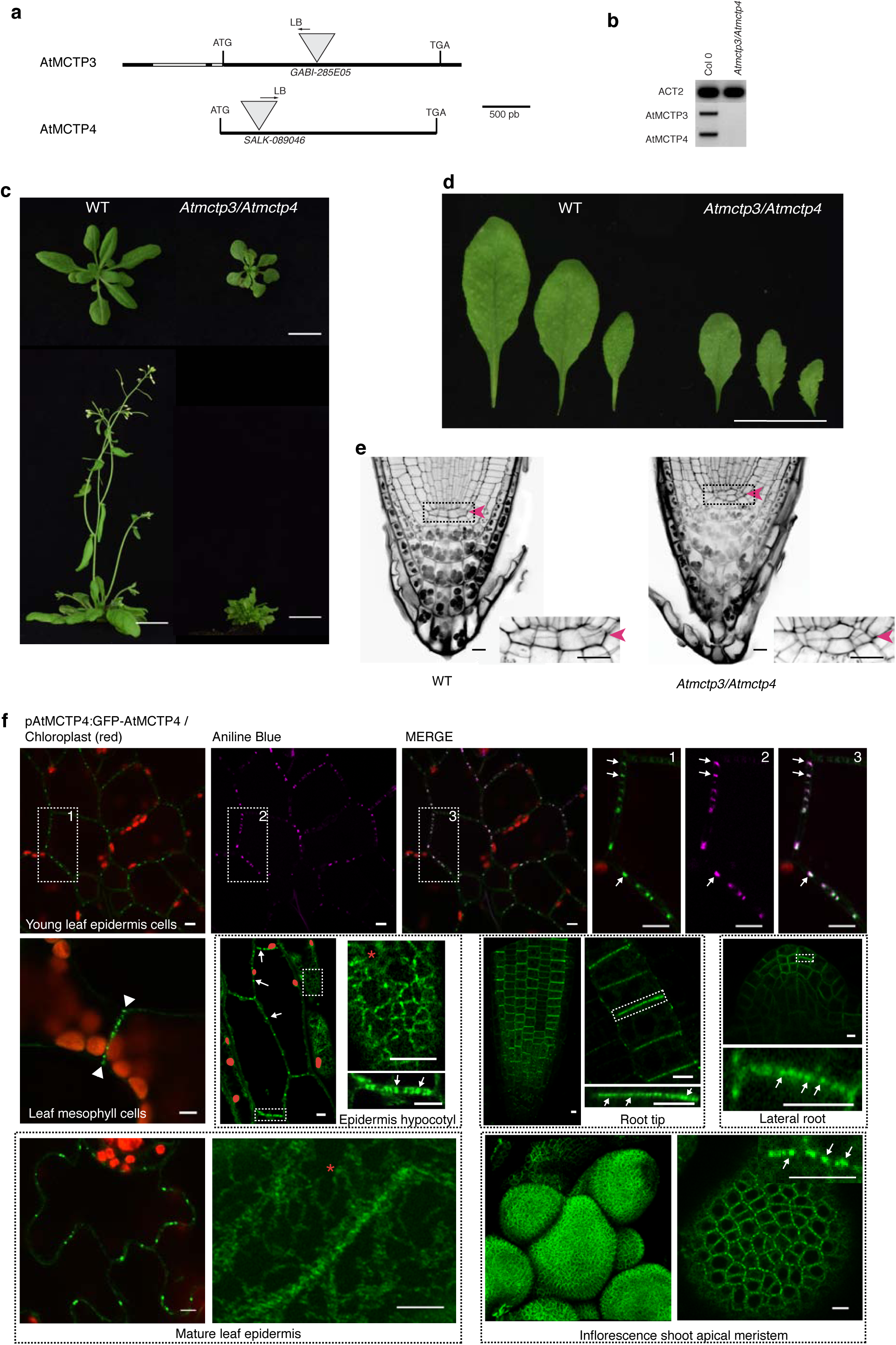
AtMCTP4 is a core plasmodesmal protein and *Atmctp3/Atmctp4* loss-of-function double mutant shows severe defects in development. **a-e**, Characterisation of *Atmctp3/Atmctp4* double mutant in *Arabidopsis*. **a**, Schematic representation of T-DNA insertions in *AtMCTP3* and *AtMCTP4*. LB, left border. **b**, RT-PCR analysis of AtMCTP3, AtMCTP4 and Actin2 (ACT2) transcripts in Col-0 wild type (WT) and *Atmctp3/Atmctp4* double mutant showing the absence of full-length transcripts in the *Atmctp3/Atmctp4* double mutant. **c**, Rosette and inflorescence stage phenotypes of *Atmctp3/Atmctp4* double mutant compared to Col-0 WT. Scale bar, 2 cm **d**, Leaf phenotypes of *Atmctp3/Atmctp4* double mutant compared to WT. Scale bar, 2 cm. **e**, Pseudo-Schiff-Propidium iodide method-stained root tips of WT and *Atmctp3/Atmctp4* double mutant. Defect in quiescent center (QC, red arrowheads) cell organisation was observed in 20 out of 20 plants examined. Scale bars, 10 μm. **f**, Subcellular localisation of GFP-AtMCTP4 under MCTP4 native promoter in *Arabidopsis* transgenic lines visualised by confocal microscopy. In all tissues examined GFP-MCTP4 shows a typical punctate distribution of plasmodesmata at the cell boundaries (indicated by white arrows). In leaf spongy mesophyll GFP-MCTP4 punctate pattern was visible only on adjoining walls (arrowheads), which contain plasmodesmata but absent from non-adjoining walls. GFP-MCTP4 dots at the cell periphery are immobile (see Suppl movie2) and co-localise perfectly with aniline blue (top row) confirming plasmodesmata localisation. In most tissues examined an ER-reticulated pattern was also observable (red stars). Boxed regions are magnified in adjacent panels. Please note that optical scan of epidermis hypocotyl were imaged in the airy scan mode and chloroplasts were manually outlined in red. Scale bars, 5 μm.

AtMCTP4 has recently been reported as an endosomal-localised protein (Liu *et al*, 2018), which is in conflict with our data indicating plasmodesmata association. To further check AtMCTP4 localisation, we expressed the protein as a GFP N-terminal fusion protein under its own promoter and analysed its localisation in *Arabidopsis* stable lines. Similar to transient expression experiments (Fig. 1), we found that pMCTP4:GFP-AtMCTP4 was located at stable punctate spots at the cell periphery (Fig. 3f white arrows; Suppl movie 2), in all tissues examined, *i.e.* leaf epidermal and spongy mesophyll cells, hypocotyl epidermis, lateral root primordia, root tip, and inflorescence shoot apical meristem. These immobile dots colocalised perfectly with aniline blue indicating plasmodesmata association (Fig.3f top row), which was also evident in leaf spongy mesophyll cells where the dotty pattern of pMCTP4:GFP-AtMCTP4 was present on adjoining walls (containing plasmodesmata), but absent from non-adjoining walls (without plasmodesmata) (Fig.3f white arrowheads). We also observed a weak but consistent ER-association of AtMCTP4 (Fig.3f, red stars).

In summary we concluded that whatever the tissue and organ considered, AtMCTP4 is strongly and consistently associated with plasmodesmata but also presents a steady association with the ER and that *Atmctp3/Atmctp4* loss-of-function is detrimental for normal plant development, including previously uncovered defect in the root apical meristem.

### The C-terminal transmembrane regions of MCTPs serve as ER-anchors

A requirement for tethers is that they physically bridge two membranes. Often this is achieved through lipid-binding module(s) at one terminus of the protein, and transmembrane domain(s) at the other (Eisenberg-Bord *et al*, 2016; Henne *et al*, 2015). All sixteen *Arabidopsis* MCTPs contain two to three predicted TMDs near their C-terminus (collectively referred to as the transmembrane region, TMR). To test whether the MCTP TMRs are determinants of ER-insertion, we generated truncation mutants lacking the C2 domains for NbMCTP7, AtMCTP3, AtMCTP4, AtMCTP6, AtMCTP9 as well as AtMCTP1/FTIP and AtMCTP15/QKY (Fig. 4a). When fused to YFP at their N-terminus, all truncated mutants retained ER-association, as demonstrated by co-localisation with RFP-HDEL (Fig. 4b left panels). Meanwhile plasmodesmata association was completely lost and the plasmodesmata-index of all truncated MCTP_TMRs dropped below one, comparable to RFP-HDEL (Fig. 4b right panels and c), quantitatively confirming the loss of plasmodesmata association when the C2 modules were deleted. We therefore concluded that, similar to the HsE-Syt and AtSYT ER-PM tether families (Giordano *et al*, 2013; Levy *et al*, 2015; Pérez-Sancho *et al*, 2015b), MCTPs insert into the ER through their TMRs, but the TMR is not sufficient for MCTP plasmodesmal localisation.

**Figure 4.**
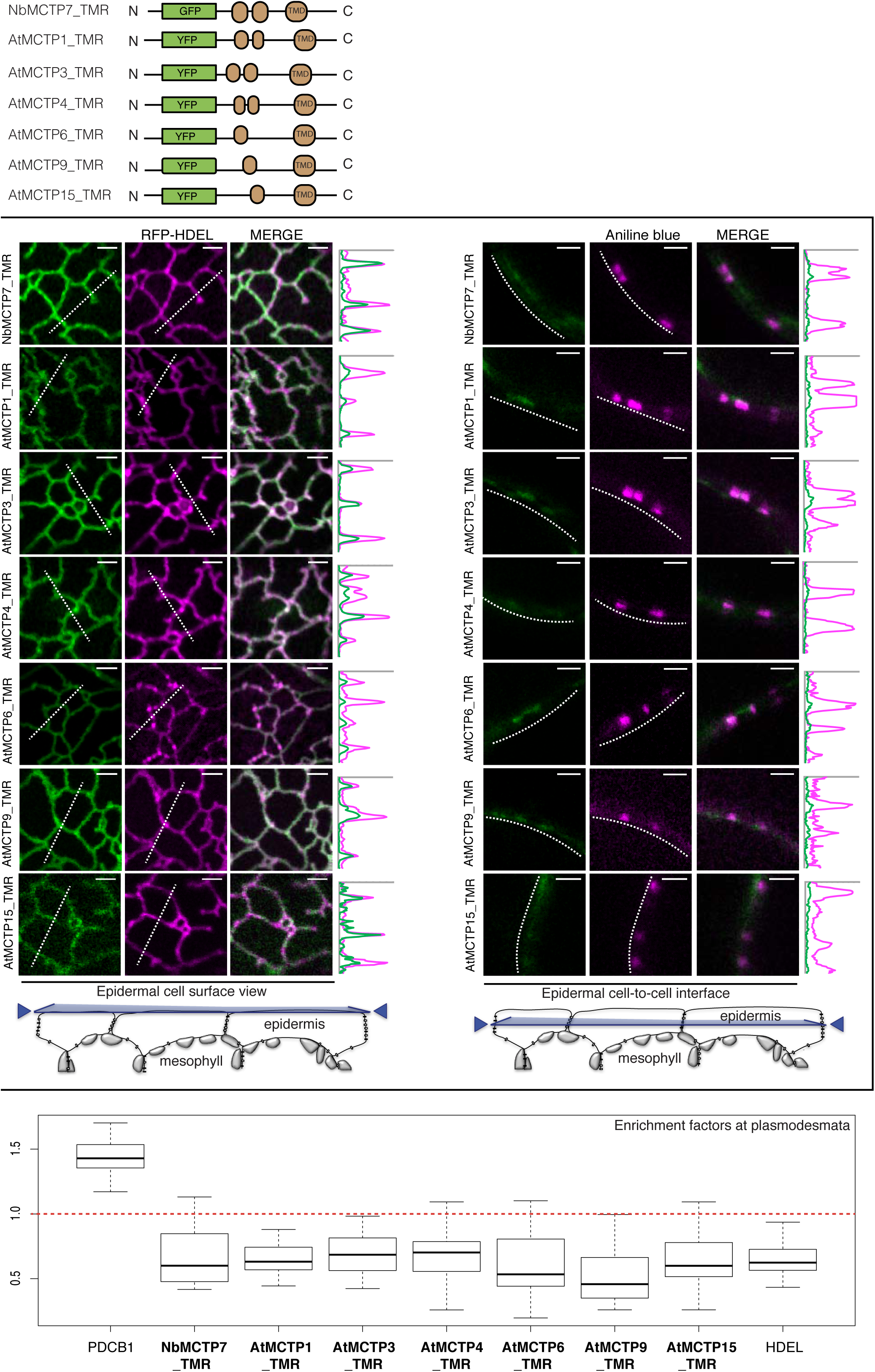
MCTPs insert into the ER membrane *via* their C-terminal transmembrane region. Localisation of truncated AtMCTPl, 3, 4, 6, 9, 15 and NbMCTP7 transmembrane regions (TMR) in *N. benthamiana* leaf epidermal cells. TMRs were tagged at their N-terminus with GFP/YFP and expressed transiently under moderate UBIQUITIN10 promoter. **a**, Schematic representation of truncated MCTPs tagged with GFP/YFP. **b**, Optical sections at cell surface (left) and cell-to-cell interface (right) showing the co-localisation between GFP/YFP-MCTP_TMR constructs and the ER-marker RFP-HDEL (left) and the plasmodesmata marker aniline blue (right). Intensity plots along the white dashed lines are shown for each colocalisation pattern. When expressed in epidermal cells, GFP/YFP-MCTP_TMR constructs associate with the ER but plasmodesmata association is lost. Scale bars, 2 μm. c, The PD index of individual truncated MCTP_TMR constructs is below 1 (red dashed line), similar to the ER marker RFP-HDEL confirming loss of plasmodesmata localisation.

### MCTP C2 domains can bind membranes in an anionic lipid-dependent manner

Members of the HsE-Syt and AtSYT tether families bridge across the intermembrane gap and dock to the PM via their C2 domains (Pérez-Sancho *et al*, 2015b; Pérez-sancho *et al*, 2016; Giordano *et al*, 2013; Saheki *et al*, 2016). *Arabidopsis* MCTPs contain three to four C2 domains, which may also drive PM-association through interactions with membrane lipids. C2 domains are independently folded structural and functional modules with diverse modes of action, including membrane docking, protein-protein interactions and calcium sensing (Corbalan-Garcia & Gómez-Fernández, 2014).

To investigate the function of MCTP C2 modules, we first searched for homologs of AtMCTP individual C2 domains (A, B, C, and D) amongst all human and *A. thaliana* proteins using the HHpred webserver (Zimmermann *et al*, 2018) for remote homology detection. The searches yielded a total of 1790 sequence matches, which contained almost all human and *A. thaliana* C2 domains. We next clustered the obtained sequences based on their all-against-all pairwise similarities in CLANS (Frickey & Lupas, 2018). In the resulting map (Supplementary Fig. 7a), the C2 domains of *Arabidopsis* MCTPs (AtMCTPs, coloured cyan) most closely match the C2 domains of membrane-trafficking and -tethering proteins, including human MCTPs (HsMCTPs, green), human Synaptotagmins (HsSyts, orange), human Ferlins (HsFerlins, blue), human HsE-Syts (HsE-Syts, magenta) and *Arabidopsis* SYTs (AtSYTs, red), most of which dock to membranes through direct interaction with anionic lipids (Giordano *et al*, 2013; Saheki *et al*, 2016; Pérez-Lara *et al*, 2016; Abdullah *et al*, 2014; Marty *et al*, 2014). By comparison to the C2 domains of these membrane-trafficking and -tethering proteins, the C2 domains of most other proteins do not make any connections to the C2 domains of AtMCTPs at the *P-value* cut-off chosen for clustering (1e-10). Thus, based on sequence similarity, the plant AtMCTP C2 domains are expected to bind membranes.

We next asked whether the C2 modules of MCTPs are sufficient for PM association *in vivo*. Fluorescent protein fusions of the C2A-D or C2B-D modules without the TMR were generated for NbMCTP7, AtMCTP3, AtMCTP4, AtMCTP6, AtMCTP9 as well as AtMCTP1/FTIP and AtMCTP15/QKY and expressed in *N. benthamiana*. We observed a wide range of sub-cellular localisations from cytosolic to PM-associated and in all cases plasmodesmata association was lost (Supplementary Fig. 7b-d).

To further investigate the potential for MCTP C2 domains to interact with membranes, we employed molecular dynamics modelling. We focussed on AtMCTP4, as a major plasmodesmal constituent and whose loss-of-function in conjunction with AtMCTP3, induces severe plant development defects (Liu *et al*, 2018) (Fig. 3). We first generated the 3D structures of all three C2 domains of AtMCTP4 using 3D homology modelling, and then tested the capacity of individual C2 to dock to membrane bilayers using coarse-grained dynamic simulations (Fig. 5a; Suppl movie 3). Molecular dynamics modelling was performed on three different membranes; 1) a neutral membrane composed of phosphatidylcholine (PC), 2) a membrane with higher negative charge composed of PC and phosphatidylserine (PS; 3:1) and 3) a PM-mimicking lipid bilayer, containing PC, PS, sitosterol and the anionic phosphoinositide phosphatidyl inositol-4-phosphate (PI4P; 57:19:20:4). The simulations showed that all individual C2 domains of AtMCTP4 can interact with lipids and dock on the membrane surface when a “PM-like” lipid composition was used (Fig. 5a). The PC-only membrane showed only weak interactions, whilst the PC:PS membrane allowed only partial docking (Fig. 5a). Docking of AtMCTP4 C2 domains arose mainly through electrostatic interactions between lipid polar heads and basic amino acid residues at the protein surface. We further tested two other MCTP members, namely AtMCTP15/QKY and NbMCTP7, which possess four rather than three C2 domains. We found that similar to AtMCTP4, the individual C2 domains of AtMCTP15/QKY and NbMCTP7 exhibited membrane interaction in the presence of the negatively charged lipids (Supplementary Fig. 8).

**Figure 5.**
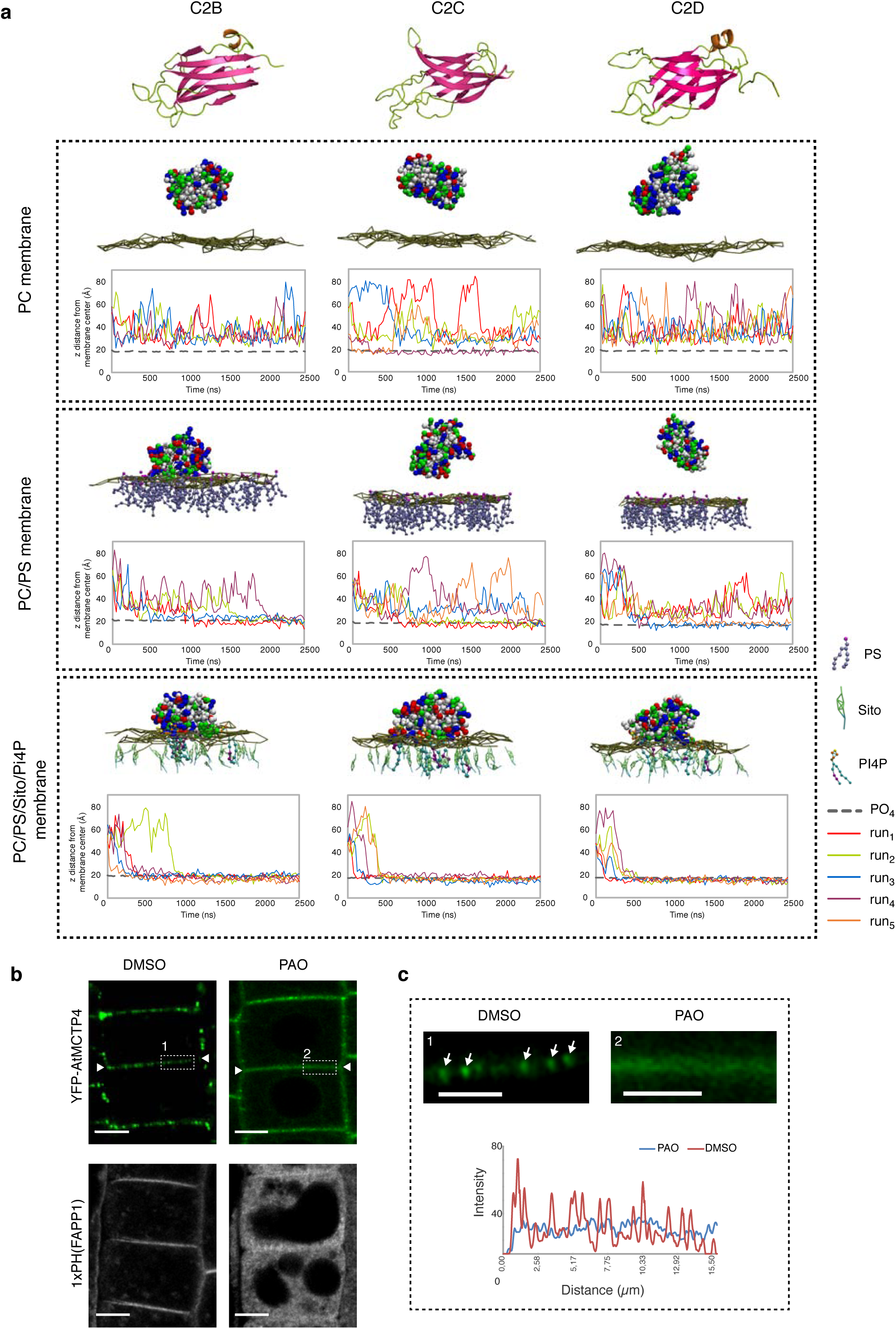
Anionic lipid-dependant membrane docking of AtMCTP4 C2 domains. **a**, Top: 3D-atomistic model of the individual AtMCTP4 C2 domains. Beta strands are shown in pink, loops in green and alpha helices in orange. Bottom: molecular dynamics of individual AtMCTP4 C2 domains with different biomimetic lipid bilayer compositions: phosphatidylcholine (PC) alone, with phosphatidylserine (PS)(PC/PS 3:1), and with PS, sitosterol (Sito) and phosphoinositol-4-phosphate (PI4P)(PC/PS/Sito/PI4P 57:19:20:4). The plots show the distance between the protein’s closest residue to the membrane and the membrane center, over time. The membrane’s phosphate plane is represented by a PO_4_ grey line on the graphs and a dark green meshwork on the simulation image captures (above graphs). For individual C2 domain and a given lipid composition, the simulations were repeated four to five times (runs 1-5). C2 membrane docking was only considered as positive when a minimum of four independent repetitions showed similarly stable interaction with the membrane. All C2 domains of AtMCTP4 show membrane interaction when anionic lipid, and in particular PI4P, are present. The amino acid colour code is as follow: red, negatively charged (acidic) residues; blue, positively charged (basic) residues; green, polar uncharged residues; and white, hydrophobic residues. **b-c**, Confocal microscopy of *Arabidopsis* root epidermal cells of UB10:YFP-AtMCTP4 transgenic lines after 40 min treatment with DMSO (mock) and PAO (60 μm), an inhibitor of PI4 kinase. To confirm PI4P depletion upon PAO treatment we used the PI4P *Arabidopsis* sensor line 1xPH(FAPP1)(Simon *et al*, 2016). PAO treatment leads to a loss of plasmodesmal punctate signal at the cell periphery (apical-basal boundary is highlighted by white arrowheads in b) for YFP-AtMCTP4, and redistribution of PM-localised 1xPH(FAPP1) to the cytoplasm (b). c, Magnified boxed-regions from b and profile plot along the cell wall after DMSO (1) or PAO (2) treatment, respectively (arrows: plasmodesmal punctae). Scale bars, 5 μm in b and 2.5μm in c.

Our molecular dynamics data thus suggests that membrane docking of the AtMCTP4 C2 domains depends on the electrostatic charge of the membrane and more specifically on the presence of PI4P, a negatively-charged lipid which has been reported as controlling the electrostatic field of the PM in plants (Simon *et al*, 2016).

To confirm the importance of PI4P for MCTP membrane interactions and thus, potentially subcellular localisation, we used a short-term treatment with phenylarsine oxide (PAO), an inhibitor of PI4-kinases (Simon *et al*, 2016). We focused on *Arabidopsis* root tips where effects of PAO have been thoroughly characterised (Simon *et al*, 2016). In control-treated roots of *Arabidopsis* plants stably expressing UB10:YFP-AtMCTP4, the fluorescent signal was most prominent at the apical-basal division plane of epidermal root cells, where numerous plasmodesmata are established during cytokinesis (Grison *et al*, 2015) (Fig. 5b white arrowheads). The YFP-AtMCTP4 fluorescence pattern was punctate at the cell periphery, each spot of fluorescence corresponding to a single or group of plasmodesmata (Fig. 5c, white arrows). We found that 40 min treatment with PAO (60 μM) induced a loss in the typical spotty plasmodesmata-associated pattern and instead AtMCTP4 became more homogenously distributed along the cell periphery (Fig.5b-c). To confirm the effect of PAO on the cellular PI4P pool, we used a PI4P-biosensor (1XPH FAPP1) which showed a clear shift from PM-association to cytosolic localisation upon treatment with PAO (Simon *et al*, 2016) (Fig. 5b). This control not only demonstrates that the PAO treatment was successful, but also highlights that the majority of PI4P was normally found at the PM, rather than the ER, of *Arabidopsis* root cells. Therefore, the effect of PAO on YFP-AtMCTP4 localisation is likely related to a perturbance of PM docking by the MCTP4 C2 domains.

Altogether, our data suggest that the C2 domains of plant MCTPs can dock to membranes in the presence of negatively charged phospholipids, and that PI4P depletion reduced AtMCTP4 stable association with plasmodesmata.

### AtMCTP4 expression is sufficient to partially restore ER-PM contacts in yeast

To further test the ability of MCTPs to physically bridge across membranes and tether the ER to the PM, we used a yeast Δtether mutant line deleted in six ER-PM tethering proteins resulting in the separation of the cortical ER (cER) from the PM (Manford *et al*, 2012) and expressed untagged AtMCTP4. To monitor recovery in cortical ER, and hence, ER-PM contacts, upon AtMCTP4 expression, we used Sec63-RFP (Metzger *et al*, 2008) as an ER marker combined with confocal microscopy. In wild-type cells, the ER was organised into nuclear (nER) and cER. The cER was visible as a thread of fluorescence along the cell periphery, covering a large proportion of the cell circumference (Fig. 6a white arrows). By contrast and as previously reported (Manford *et al*, 2012), we observed a substantial reduction of cER in the Δtether mutant, with large areas of the cell periphery showing virtually no associated Sec63-RFP (Fig. 6a). When AtMCTP4 was expressed into the Δtether mutant line, we observed partial recovery of cER, visible as small regions of Sec63-RFP closely apposed to the cell cortex. We further quantified the extent of cER in the different lines by measuring the ratio of the length of cER (Sec63-RFP) against the cell perimeter (through calcofluor wall staining) and confirmed that ATMCTP4 expression induced an increase of cER from 7.3 % to 23.1% when compared to the Δtether mutant (Fig. 6b). This partial complementation is similar to results obtained with yeast deletion mutants containing only a single endogenous ER-PM tether, IST2, or all three isoforms of the tricalbin (yeast homologs of HsE-Syts) (Manford *et al*, 2012), supporting a role of AtMCTP4 as ER-PM tether.

**Figure 6.**
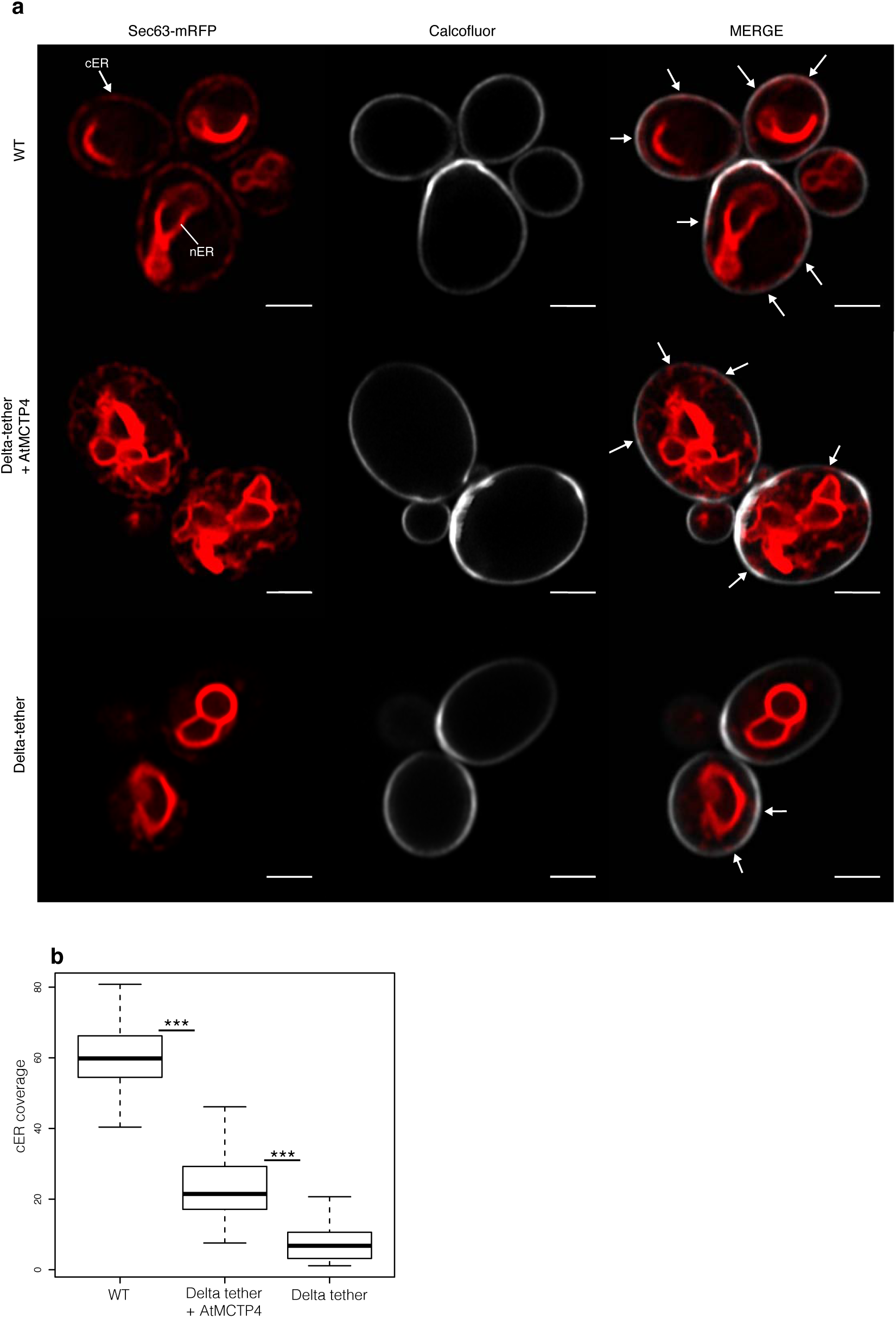
AtMCTP4 expression in yeast partially restores ER-PM membrane contact sites. Expression of AtMCTP4 in yeast Δtether cells (ist2Δ, scs2/22Δ, and tcb1/2/3Δ) (Manford *et al*, 2012) followed by confocal microscopic analysis of cortical ER. **a**, Top to bottom: Wild type (WT) cell, Δtether expressing untagged AtMCTP4 and Δtether cells, respectively. The cortical ER (cER) and nuclear ER (nER) are labelled by the ER marker Sec63-RFP (red), while the cell periphery is stained by calcofluor (white). In WT cells, both nER and cER are visible, whereas in Δtether cell only remains of the cER are visible (arrows), due to the loss of ER-PM tethering factors. When AtMCTP4 is expressed in the yeast Δtether, partial recovery of cER is observable (arrows). Scale bars, 2 μm. **b**, Quantification of cER expressed as a ratio of the length of cER to length of the PM in WT, Δtether+AtMCTP4 and Δtether cells. Numbers of cells used for quantifying the cER: n = 39 for WT, n=49 for Δtether+AtMCTP4 and n=61 for Δtether strains. Wilcoxon test was used to compare the extent of cER between the different strains i.e. WT *versus* Δtether+AtMCTP4 and Δtether+AtMCTP4 *versus* Δtether (\*\*\**p-value* < 0.001).

## Discussion

In plants, communication between cells is facilitated and regulated by plasmodesmata, ~50 nm diameter pores that span the cell wall and provide cell-to-cell continuity of three different organelles: the PM, cytoplasm, and ER. The intercellular continuity of the ER and the resulting architecture of the pores make them unique amongst eukaryotic cellular junctions, and qualify plasmodesmata as a specialised type of ER-PM MCS (Bayer *et al*, 2017; Tilsner *et al*, 2016). Like other types of MCS, the membranes within plasmodesmata are physically connected but so far the molecular components and function of the ER-PM tethering machinery remain an enigma.

Here, we provide evidence that members of the MCTP family, some of which have been described as key regulators of intercellular trafficking and cell-to-cell signalling (Vaddepalli *et al*, 2014; Liu *et al*, 2018, 2012), also act as ER-PM tethers inside the plasmodesmata pores.

### MCTPs as core plasmodesmal components

Whi1st several MCTPs have previously been characterised as regulators of cell-to-cell trafficking or signalling (Liu *et al*, 2012, 2018; Vaddepalli *et al*, 2014; Liu *et al*, 2017), only some have also been localised to plasmodesmata, Whilst other studies reported alternative localisations which include PM, ER, Golgi, endosomes and cytoplasm (Trehin *et al*, 2013; Liu *et al*, 2017, 2018, 2012; Kraner *et al*, 2017; Vaddepalli *et al*, 2014), perhaps depending on the isoform, orientation of fluorescent protein fusions and expression context. Here, we have identified several MCTPs (6-10 out of 16) as belonging to the most abundant proteins at plasmodesmata through both *in vivo* and proteomic data. This includes AtMCTP3 and AtMCTP4 recently identified as modulators of SHOOTMERISTEMLESS trafficking (Liu *et al*, 2018), for which we find that a *Atmctp3/Atmctp4* loss-of-function mutant displays a severe developmental phenotype, including defects in the root QC, that agrees with the findings of Liu *et al*, 2018. Whereas Liu *et al*. (2018) observed endosomal-localisation of AtMCTP3 and AtMCTP4, our data suggest they are primarily plasmodesmata-associated. We therefore propose that MCTPs are core plasmodesmata-constituents and that AtMCTP3 and AtMCTP4 may possibly regulate the transport of SHOOTMERISTEMLESS, at the pores.

### MCTPs as plasmodesmata-specific ER-PM tethers

While ER-PM contacts within plasmodesmata have been observed for decades (Ding *et al*, 1992; Tilsner *et al*, 2011; Tilney *et al*, 1991; Nicolas *et al*, 2017b), the molecular identity of the tethers has remained elusive. Here we propose that MCTPs are prime plasmodesmal membrane tethering candidates as they possess all required features: 1) strong association with plasmodesmata; 2) structural similarity to known ER-PM tethers such as HsE-Syts and AtSYTs (Levy *et al*, 2015; Pérez-Sancho *et al*, 2015b; Giordano *et al*, 2013) possessing an ER-inserted TMR at one end and multiple lipid-binding C2 domains at the other for PM docking; 3) the ability to partially restore ER-PM tethering in a yeast Δtether mutant.

Similarly to other ER-PM tethers (Eisenberg-Bord *et al*, 2016; Wong *et al*, 2016; Henne *et al*, 2015; Giordano *et al*, 2013), MCTP C2 domains dock to the PM through electrostatic interaction with anionic lipids, especially PI4P and to a lesser extent PS. In contrast with animal cells, PI4P is found predominantly at the PM in plant cells and defines its electrostatic signature (Simon *et al*, 2016). Although plasmodesmata are MCS, they are also structurally unique: both the ER and the PM display extreme, and opposing membrane curvature inside the pores; the ER tubule is linked to the PM on all sides; and the membrane apposition is unusually close (2-3 nm in type I post-cytokinetic pores (Nicolas *et al*, 2017a)). So while structurally related to known tethers, MCTPs are also expected to present singular properties. For instance, similar to the human MCTP2, MCTPs could favour ER membrane curvature through their TMR (Joshi *et al*, 2017). Plasmodesmata also constitute a very confined environment, which together with the strong negative curvature of the PM, may require the properties of MCTP C2 domains to differ from that of HsE-Syts or AtSYTs. All of these aspects will need to be investigated in the future.

### Inter-organellar signalling at the plasmodesmal MCS?

In yeast and animals, MCS have been shown to be privileged sites for inter-organelle signalling by promoting fast, non-vesicular transfer of molecules such as lipids (Gallo *et al*, 2016; Saheki *et al*, 2016; Wong *et al*, 2016). Unlike the structurally analogous tethering proteins AtSYTs and HsE-Syts, MCTPs do not harbour known lipid-binding domains that would suggest that they participate directly in lipid transfer between membranes. However, MCTPs are likely to act in complex with other proteins (Fulton *et al*, 2009; Trehin *et al*, 2013) which may include lipid shuttling proteins. For instance, AtSYT1, which contains a lipid-shuttling SMP (synaptotagmin-like mitochondrial-lipid binding protein) domain (Reinisch & De Camilli, 2016), is recruited to plasmodesmata during virus infection and promotes virus cell-to-cell movement (Levy *et al*, 2015). MCS tethers typically interact with other MCS components and locally regulate their activity, act as Ca^2+^ sensors, or modulate membrane spacing to turn lipid shuttling on or off (Eden *et al*, 2010, 2016; Ho *et al*, 2016; Chang *et al*, 2013; Kim *et al*, 2015; Giordano *et al*, 2013; Idevall-Hagren *et al*, 2015a; Petkovic *et al*, 2014; Zhang *et al*, 2005; Fernàndez-Busnadiego *et al*, 2015; Omnus *et al*, 2016; Saheki *et al*, 2016). Similar activities could be performed by MCTPs at plasmodesmata. To date however, ER-PM cross-talk at plasmodesmata remains hypothetical.

### Combining organelle tethering and cell-to-cell signalling functions

Several members of the MCTP family have previously been implicated in regulating either macromolecular trafficking or intercellular signalling through plasmodesmata. AtMCTP1/FTIP interacts with, and is required for phloem entry of the Flowering Locus T (FT) protein, triggering transition to flowering at the shoot apical meristem (Liu *et al*, 2012). Similarly, AtMCTP3/AtMCTP4 regulate trafficking of SHOOTMERISTEMLESS in the shoot apical meristem, however in this case they prevent cell-to-cell trafficking (Liu *et al*, 2018). AtMCTP15/QKY promotes the transmission of an unidentified non-cell-autonomous signal through interaction with the plasmodesmata/PM-located receptor-like kinase STRUBBELIG (Vaddepalli *et al*, 2014). Thus, previously characterised MCTP proteins regulate intercellular trafficking/signalling either positively or negatively.

Whilst the mechanisms by which these MCTP proteins regulate intercellular transport/signalling have not been elucidated, MCTP physical interaction with mobile factors or receptor is critical for proper function (Vaddepalli *et al*, 2014; Liu *et al*, 2017, 2018, 2012). In AtMCTP1/FTIP, the interaction is mediated by the C2 domain closest to the TMR (Liu *et al*, 2017). For the C2 domains of HsE-Syts, conditional membrane docking is critical for their function and depends on intramolecular interactions, cytosolic Ca^2+^ and the presence of anionic lipids (Idevall-Hagren *et al*, 2015b; Fernàndez-Busnadiego *et al*, 2015; Saheki *et al*, 2016; Bian *et al*, 2018; Giordano *et al*, 2013). With three to four C2 domains, it is conceivable that MCTPs assume different conformations within the cytoplasmic sleeve in response to changes in the plasmodesmal PM composition, Ca^2+^, and the presence of interacting mobile signals (Fig.7), which could link membrane tethering to cell-cell signalling. Understanding in detail how MCTPs function in the formation and regulation of the plasmodesmal MCS will be an area of intense research in the coming years.

**Figure 7.**
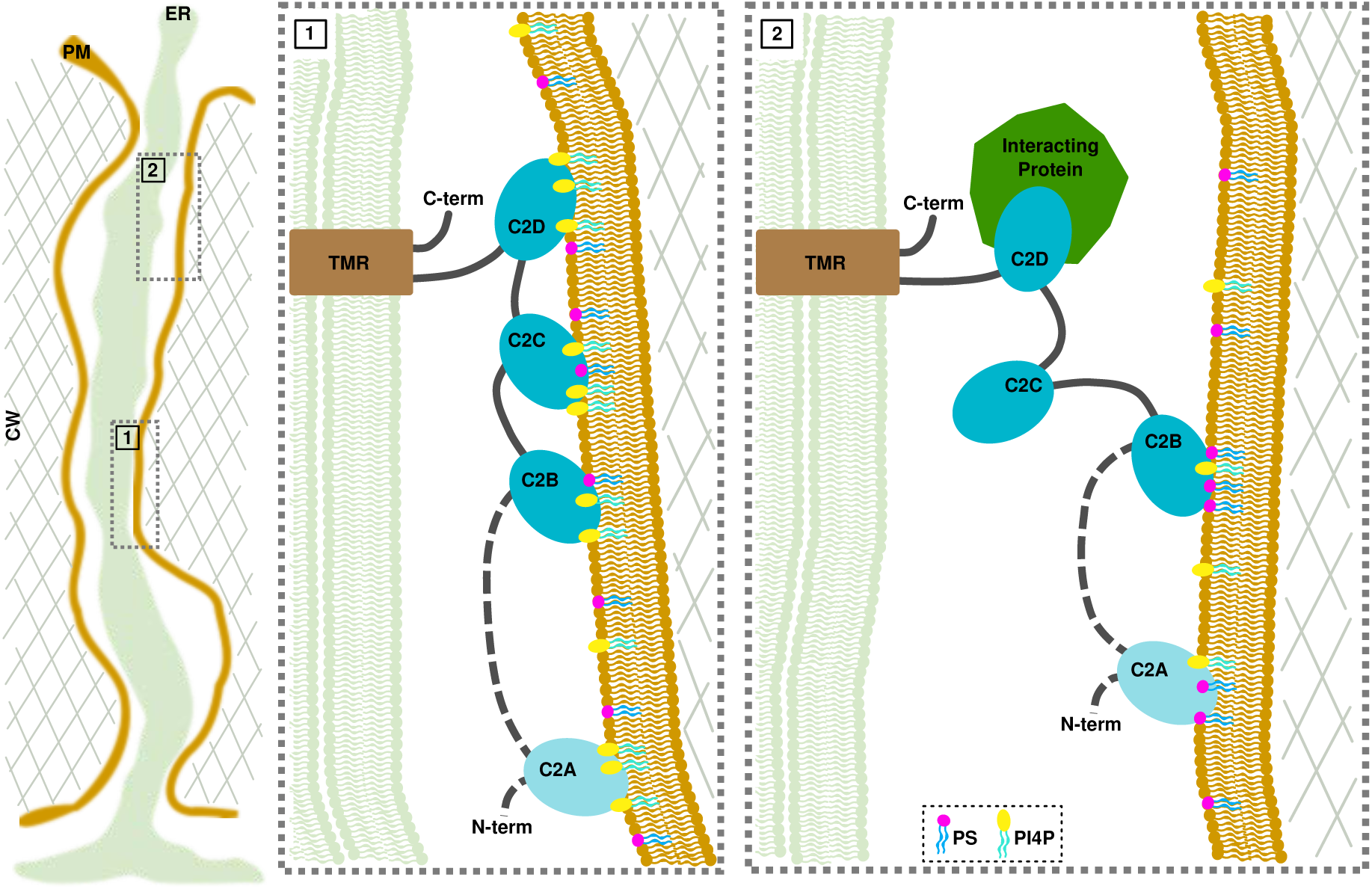
Model of MCTP arrangement within plasmodesmata and hypothetical conditional docking events. Inside plasmodesmata, MCTPs insert into the ER *via* their transmembrane regions (TMR), while docking to the PM by interacting with the negatively-charged phospholipids, PS and PI4P *via* their C2 domains. In condition of high PI4P/PS levels, all C2 domains interact with the PM, maintaining the ER close to the PM (panel 1). Decrease in the PI4P pool and/or protein interaction causes a detachment of some but not all C2 domains, which then modulate the space between the two membranes and the properties of the cytoplasmic sleeve. Please note that the exact topology of the TMR is not currently known.

## Material & Methods

### Biological material and growth conditions

*Arabidopsis* (*Columbia*) and transgenic lines were grown vertically on solid medium composed of *Murashige and Skoog* (MS) medium including vitamins (2.15g/L), MES (0.5g/L) and plant-Agar (7g/L), pH 5.7, then transferred to soil under long-day conditions at 22 °C and 70% humidity.

*Arabidopsis (Landsberg erecta)* culture cells were cultivated as described in(Nicolas *et al*, 2017a) under constant light (20μE/m/s) at 22°C. Cells were used for experimentation at various ages ranging from four to seven-day-old (mentioned in individual experiment).

### MCTP sequence alignment and phylogenetic tree

The 16 members of *Arabidopsis thaliana* MCTP family, gathering a total of 59 C2 domains, were dissected using a combination of several bioinformatic tools. The alignment of *A. thaliana* MCTP members from(Liu *et al*, 2017) combined with Pfam predictions was used as a first step to segregate the MCTP members into “sub-families”: the short MCTPs, which contain three C2 domains (C2B to C2D) and the long MCTPs, which contain four C2 domains (C2A to C2D). The short MCTPs lack the C2A domain, whereas the C2B-C-D are conserved in all members.

The prediction and delimitation of C2 domains in proteins, including MCTPs, from databases such as Pfam are rather imprecise. In order to provide stronger and more accurate predictions for the delimitation of each C2 domain, we used both the PSIRED(Buchan *et al*, 2013; Jones, 1999) protein sequence analysis (http://bioinf.cs.ucl.ac.uk/psipred/) and Hydrophobic Cluster Analysis(Callebaut *et al*, 1997) (HCA; http://www-ext.impmc.upmc.fr/~callebau/HCA.html). Multiple sequence alignment was performed using Clustal Omega (http://www.ebi.ac.uk/Tools/msa/clustalo/).

### Cluster map of Human and *A. thaliana* C2 domains

To generate a C2 cluster map, we first collected all *A. thaliana* and human C2 domains, using the HHpred webserver(Alva *et al*, 2016; Söding *et al*, 2005). The obtained set was filtered to a maximum of 100% pairwise sequence identity at a length coverage of 70% using MMseqs2(Steinegger & Söding, 2017) to eliminate all redundant sequences. The sequences in the filtered set, comprising almost all human and *A. thaliana* C2 domains (~1800 in total), was next clustered in CLANS(Frickey & Lupas, 2018) based on their all-against-all pairwise sequence similarities as evaluated by BLAST P-values.

### Cloning of MCTPs and transformation into *Arabidopsis*

The different constructs used in this study were either PCR amplified from cDNA or genomic DNA (Col-0) using gene specific primers (Supplementary Table S2), or were synthesised and cloned into donor vectors by GenScript^®^ (Supplementary Table S2). For N-terminal tag fusion, the PCR/DNA products were cloned into the Multisite Gateway^®^ donor vectors pDONR-P2RP3 (Invitrogen, Carlsbad, CA), and then subcloned into pB7m34GW or pK7m34GW using the multisite LR recombination system(Karimi *et al*, 2002), the moderate promoter UBIQUITIN10 (UBQ10/pD0NR-P4P1R previously described in(Marquès-Bueno *et al*, 2016)) and eYFP/pD0NR221. For C-terminal tag fusion, the PCR/DNA products were first cloned into pD0NR221, then multisite recombined using a mVenus/pD0NR-P2RP3 and UBQ10/pD0NR-P4P1R.

For the expression of GFP-MCTP4 driven by its native promotor we used the binary vector pRBbar-OCS harboring a BASTA resistance, a multiple cloning side (MCS) and an octopine synthase (OCS) terminator within the left and right borders. The vector derived from the pB2GW7 (Karimi *et al*, 2002) by cutting out the expression cassette with the restriction enzymes SacI and HindIII and replaced it with a synthesized MCS and an OCS terminator fragment. To combine promoter region and GFP-MCTP4 coding sequence we used In-Fusion cloning (Takara Bio Europe). To PCR amplify the coding sequence for GFP-MCTP4 with its respective primers (Supplementary Table2) we used the plasmid coding for GFP-MCTP4 as template (previously described as GFP-C2-89 by (Kraner *et al*, 2017)). The resulting pRBbar-pMCTP4: plasmid was linearized with BamH1/Pst1 the amplified GFP-MCTP4 was fused in to generate the MCTP4 promoter driven GFP-MCTP4 construct (pMCTP4:GFP-MCTP4). Expression vectors were transformed in *Arabidopsis* Col-0 by floral dip(Clough & Bent, 1998), and transformed seeds were selected based on plasmid resistance.

*N. benthamiana* homologs of Arabidopsis MCTP isoforms were identified by protein BLAST searches against the SolGenomics *N. benthamiana* genome (https://solgenomics.net). An ortholog of AtMCTP7, NbMCTP7 (Niben101Scf03374g08020.1) was amplified from *N. benthamiana* leaf cDNA. The recovered cDNA of NbMCTP7 differed from the SolGenomics reference by the point mutation G287D and three additional silent nucleotide exchanges, as well as missing base pairs 1678-1716 which correspond to thirteen in-frame codons (encoding the amino acid sequence LKKEKFSSRLHLR). We note that this nucleotide and amino acid sequence is exactly repeated directly upstream (bp 1639-1677) in the SolGenomics reference and may thus represent an error in the *N. benthamiana* genome assembly. The recovered NbMCTP7 sequence has been submitted to database.

### Generation of *Atmctp3/Atmctp4* loss-of-function Arabidopsis mutant

*Atmctp3* (Sail_755_G08) and *Atmctp4* (Salk_089046) T-DNA insertional Arabidopsis mutants (background Col-0) were obtained from the Arabidopsis Biological Resource Center (http://www.arabidopsis.org/). Single T-DNA insertion lines were genotyped and homozygous lines were crossed to obtain double homozygous *Atmctp3/Atmctp4*.

For genotyping, genomic DNA was extracted from Col-0, *Atmctp3* (GABI-285E05) and *Atmctp4* (SALK-089046) plants using chloroform:isoamyl alcohol (ratio24:1), genomic DNA isolation buffer (200mM Tris HCL PH7.5, 250mM NaCl, 25mM EDTA and 0.5% SDS) and isopropanol. PCR were performed with primers indicated in Supplementary Table2. For transcript expression, total mRNA was extracted from Col-0 and *Atmctp3/Atmctp4* using RNeasy^®^ Plant Mini Kit (QIAGEN) and cDNA was produced using random and oligodT primers. The expression level of AtMCTP3, AtMCTP4 and ubiquitous Actin2 (ACT2) transcript was tested by PCR amplification using primers listed in Supplementary Table2.

### Confocal Laser Scanning Microscopy

For transient expression in *N. benthamiana*, leaves of 3 week-old plants were pressure-infiltrated with GV3101 agrobacterium strains, previously electroporated with the relevant binary plasmids. Prior to infiltration, agrobacteria cultures were grown in Luria and Bertani medium with appropriate antibiotics at 28°C for two days then diluted to 1/10 and grown until the culture reached an OD_600_ of about 0.8. Bacteria were then pelleted and resuspended in water at a final OD_600_ of 0.3 for individual constructs, 0.2 each for the combination of two. The ectopic silencing suppressor 19k was co-infiltrated at an OD_600_ of 0.15. Agroinfiltrated *N. benthamiana* leaves were imaged 3-4 days post infiltration at room temperature. ~ 2 by 2 cm leaf pieces were removed from plants and mounted with the lower epidermis facing up onto glass microscope slides.

Transgenic *Arabidopsis* plants were grown as described above. For primary roots, lateral roots and hypocotyl imaging, six to seven days old seedlings or leaves of 5-8 leaf stage rosette plants were mounted onto microscope slides. For shoot apical meristem imaging, the plants were first dissected under a binocular then transferred to solid MS media and immediately observed using a water-immersion long-distance working 40X water immersion objective. Confocal imaging was performed on a Zeiss LSM 880 confocal laser scanning microscope equipped with fast AiryScan using Zeiss C PL APO x63 oil-immersion objective (numerical aperture 1.4). For GFP, YFP and mVenus imaging, excitation was performed with 2-8% of 488 nm laser power and fluorescence emission collected at 505-550 nm and 520-580 nm, respectively. For RFP and mCherry imaging, excitation was achieved with 2-5% of 561 nm laser power and fluorescence emission collected at 580-630 nm. For aniline blue (infiltrated at the concentration of 25 μg/mL) and Calcofluor White (1 μg /mL), excitation was achieved with 5% of 405 nm laser and fluorescence emission collected at 440-480 nm. For colocalisation sequential scanning was systematically used.

For quantification of NbMCTP7 co-localisation with VAP27.1, SYT1 and PDCB1, coexpression of the different constructs was done in *N. benthamiana*. An object based method was used for colocalization quantification(Bolte & Cordelières, 2006). Images from different conditions are all acquired with same parameters (zoom, gain, laser intensity etc.) and channels are acquired sequentially. These images are processed and filtered using ImageJ software (https://imagei.nih.gov/ij/) in order to bring out the foci of the pictures. These foci were then automatically segmented by thresholding and the segmented points from the two channels were assessed for colocalization using the ImageJ plugin *Just Another Colocalization Plugin* (*JACoP*)(Bolte & Cordelières, 2006). This whole process was automatized using a macro (available upon demand).

Pseudo-Schiff-Propidium iodide stained *Arabidopsis* root tips was performed according to(Truernit *et al*, 2008). Aniline blue staining was performed according to(Grison *et al*, 2015). Brightness and contrast were adjusted on ImageJ software (https://imagej.nih.gov/ij/).

### Plasmodesmata (PD) index

Plasmodesmata depletion or enrichment was assessed by calculating the fluorescence intensity of GFP/YFP-tagged full-length MCTP, truncated MCTPs and the proton pump ATPase GFP-PMA2(Gronnier *et al*, 2017), at 1) plasmodesmata (indicated by mCHERRY-PDCB1, PDLPl-mRFP or aniline blue) and 2) at the cell periphery (i.e. outside plasmodesmata pitfields). For that, confocal images of leaf epidermal cells (*N. benthamiana* or *Arabidopsis*) were acquired by sequential scanning of mCHERRY-PDCB, PDLP1-mRFP or aniline blue (plasmodesmata markers) in channel 1 and GFP/YFP-tagged MCTPs in channel 2 (for confocal setting see above). About thirty images of leaf epidermis cells were acquired with a minimum of three biological replicates. Individual images were then processed using ImageJ by defining five regions of interest (ROI) at plasmodesmata (using plasmodesmata marker to define the ROI in channel1) and five ROIs outside plasmodesmata. The ROI size and imaging condition were kept the same. The GFP/YFP-tagged MCTP mean intensity (channel 2) was measured for each ROI then averaged for single image. The plasmodesmata index corresponds to intensity ratio between fluorescence intensity of MCTPs at plasmodesmata versus outside the pores. For the plasmodesmata-index of RFP-HDEL, PDLP1-RFP and mCHERRY-PDCB1 we used aniline to indicate pitfields. R software was used for making the box plots and statistics.

### FRAP analysis

For FRAP analysis, GFP-NbMCTP7, RFP-HDEL and mCHERRY-PDCB1-expressing *N. benthamiana* leaves were used. The experiments were performed on a Zeiss LSM 880 confocal microscope equipped with a Zeiss C PL APO x63 oil-immersion objective (numerical aperture 1.4). GFP and mCherry were respectively excited at 488nm and 561nm with 2% of Argon or DPSS 561-10 laser power, and fluorescence was collected with the GaAsp detector at 492-569nm and 579-651nm, respectively. To reduce as much as possible scanning time during FRAP monitoring, the acquisition window was cropped to a large rectangle of 350 by 50 pixels, with a zoom of 2.7 and pixel size of 0.14μm. By this mean, pixel dwell time was of 0.99μs and total frame scan time could be reduced down to 20 ms approximately. Photobleaching was performed on rectangle ROIs for the ER-network and on circle ROIs for the pitfields with the exciting laser wavelengths set to 100%. The FRAP procedure was the following: 30 pre-bleach images, 10 iterations of bleaching with a pixel dwell time set at 1.51 μs and then 300 images post-bleach with the “safe bleach mode for GaAsp”, bringing up the scan time up to approximately 200ms. The recovery profiles were background substracted and then double normalized (according to the last prebleach image and to the reference signal, in order to account for observational photobleaching) and set to full scale (last pre-bleach set to 1 and first post-bleach image set to 0), as described by Kote Miura in his online FRAP-teaching module (EAMNET-FRAP module, https://embl.de). Plotting and curve fitting was performed on GraphPad Prism (GraphPad Software, Inc.).

### 3D-SIM imaging

For 3D structured illumination microscopy (3D-SIM), an epidermal peal was removed from a GFP-NbMCTP7-expressing leaf and mounted in Perfluorocarbon PP11(Littlejohn *et al*, 2014) under a high precision (170mm+/-5mm) coverslip (Marie Enfield). The sample chamber was sealed with non-toxic Exaktosil N 21 (Bredent, Germany). 3D-SIM images were obtained using a GE Healthcare / Applied Precision OMX v4 BLAZE with a 1.42NA Olympus PlanApo N 60X oil immersion objective. GFP was excited with a 488nm laser and imaged with emission filter 504-552nm (528/48nm). SR images were captured using Deltavison OMX software 3.70.9220.0. SR reconstruction, channel alignment and volume rendering were done using softWoRx V. 7.0.0.

### Yeast

Wild-type (SEY6210) and delta-tether yeast strain(Manford *et al*, 2012) were transformed with Sec63.mRFP (pSM1959). Sec63.mRFP(Metzger *et al*, 2008) was used as an ER marker and was a gift from Susan Mickaelis (Addgene plasmid #41837). Delta-tether/Sec63.mRFP strain was transformed with AtMCTP4 (pCU416: pCU between SacI and SpeI sites, Cyc1 terminator between XhoI and KpnI sites and AtMCTP4 CDS between BamHI and SmaI sites, Supplementary table S2). Calcofluor White was used to stain the cell wall of yeast. All fluorescent microscopy was performed on midlog cells, grown on selective yeast media (-URA -LEU for AtMCTP4 and Sec63 expression, and -LEU for Sec63). Images were acquired with Airyscan module, using a 63X oil immersion lens and sequential acquisition. Brightness and contrast were adjusted on ImageJ software (https://imagei.nih.gov/ii/).

### Supplementary methods

Methods for plasmodesmata label-free proteomic analysis and dynamic modelling are described in details in Supplementary method1.

Sequence data for genes in this article can be found in the GenBank/EMBL databases using the following accession numbers: AtMCTP1, At5g06850; AtMCTP2, At5g48060; AtMCTP3, At3g57880; AtMCTP4, At1g51570; AtMCTP5, At5g12970; AtMCTP6, At1g22610; AtMCTP7, At4g11610; AtMCTP8, At3g61300; AtMCTP9, At4g00700; AtMCTP10, At1g04150; AtMCTP11, At4g20080; AtMCTP12, At3g61720; AtMCTP13, At5g03435; AtMCTP14, At3g03680; AtMCTP15, At1g74720; AtMCTP16, At5g17980 and NbMCTP7, Niben101Scf03374g08020.1.

## Acknowledgements

This work was supported by the National Agency for Research (Grant ANR-14-CE19-0006-01 to E.M.B), the European Research Council (ERC) under the European Union’s Horizon 2020 research and innovation programme (grant agreement No 772103-BRIDGING to E. M.B), Fonds National de la Recherche Scientifique (NEAMEMB PDR T.1003.14, BRIDGING CDR J.0114.18 and RHAMEMB CDR J.0086.18 to L. L. and M D.). J.D.P. is funded by a PhD fellowship from the Belgian “Formation à la Recherche dans l’Industrie et l’Agriculture” (FRIA). Work in J.T, lab is supported by grant BB/M007200/1 from the U.K. Biotechnology and Biomedical Sciences Research Council (BBSRC).

Fluorescence microscopy analyses were performed at the plant pole of the Bordeaux Imaging Centre (http://www.bic.u-bordeaux.fr). The proteomic analyses were performed at the Functional Genomic Center of Bordeaux, (https://proteome.cgfb.u-bordeaux.fr). We thank Steffen Vanneste and Abel Rosado for providing the VAP27.1.RFP and SYT1.GFP binary vectors and Yvon Jaillais for providing the 1xPH(FAPP1) *Arabidopsis* transgenic lines. The plasmid pRBbar-OCS was kindly provided by Prof. Frederik Börnke (IGZ—Leibniz Institute of Vegetable and Ornamental Crops, Großbeeren, Germany). We thank Christophe Trehin and Patrice Morel for providing the AtMCT15_C2s construct and Alenka Copic for providing the yeast WT and Δ-tether strains. We thank Fabrice Cordelières for his help for the fluorescence image quantification and Paul Gouget, Yvon Jaillais, Andrea Paterlini, and Yrjo Helariutta for critical review of the article prior to submission.

## Contributions

F. I., M.S.G., M.F. and S.C. carried out the proteomic analysis. M.L.B. cloned the MCTPs, produced and phenotyped the *Arabidopsis* transgenic lines, with the exception of AtMCTP4:GFP-MCTP4 and 35S:GFP-MCTP6 which were generated by M.K.. M.L.B. and J.D.P. imaged the MCTP reporter lines. W.N. carried out the FRAP analysis and image quantification for co-localisation with the help of L.B.. A.G. performed the phylogenic analysis. J.D.P. carried out the PAO experiments. M.L.B. performed the yeast experiments. T.J.H. and J.T. performed the 3D-SIM. V.A. carried out the C2 cluster map analysis. J.D.P. carried out the molecular dynamic analysis with the help of J-M.C. and L.L..

E.M.B. conceived the study and designed experiments with the help of J.T and L.L. E.M.B, J.D.P., J.T. and M.L.B. wrote the manuscript. All the authors discussed the results and commented on the manuscript.

---

**Supplementary Figure 1.**
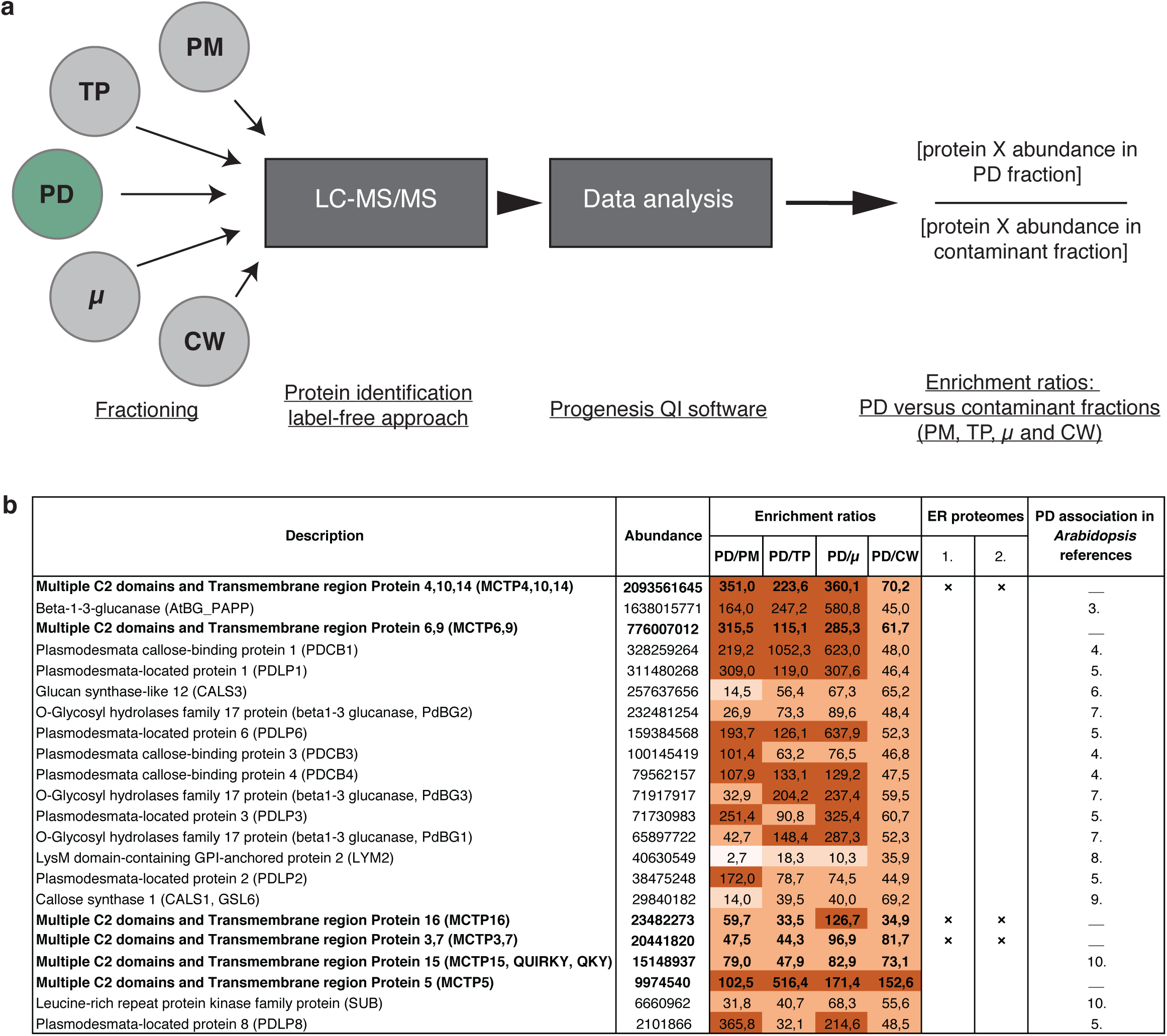
MCTP members are highly enriched in the *Arabidopsis* plasmodesmata core proteome. (a) Label-free quantitation strategy was used to determine the relative abundance of proteins in the plasmodesmata (PD) fraction *versus* contaminant subcellular fractions namely, the PM, total extract (TP), microsomes (μ) and cell wall (CW). (b) Selected set of proteins from the plasmodesmata core proteome (see Supplementary Table1 for the complete list) showing the abundance and enrichment ratios of known plasmodesmal proteins (reference to published papers is indicated below the table) and MCTP members (in bold). MCTP members are present in the plasmodesmal core proteome being both abundant and highly enriched (from 47.5- to 351-folds compared to the PM) similar to known plasmodesmata proteins. Please note that in some cases, the identified peptides did not permit unambiguous identification of MCTP isoforms due to high sequence homology between several members. The different shades (light to dark) of brown represent different enrichment levels (0-10; 10-20; 20-100 and above 100).

**Supplementary Figure 2.**
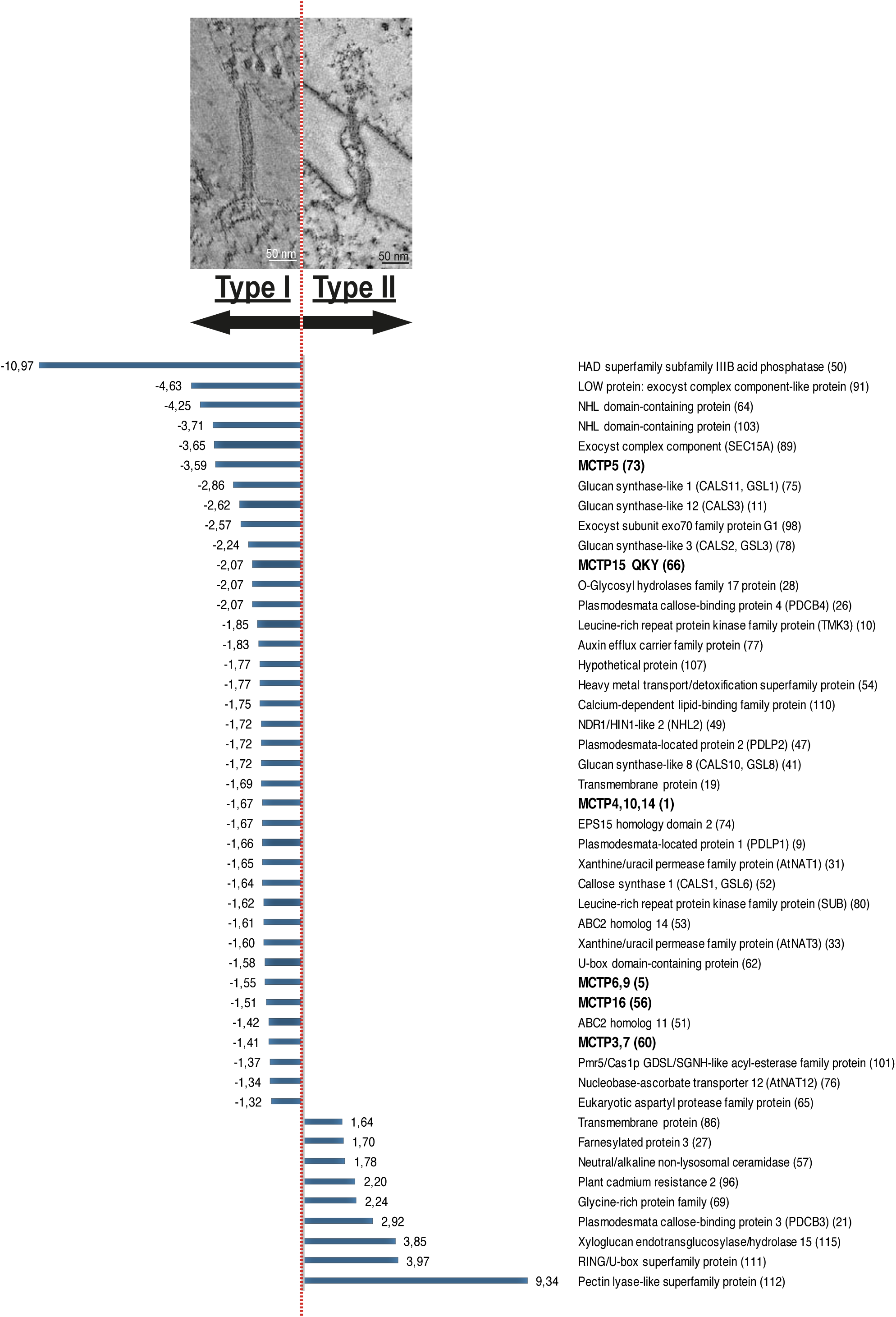
Differential abundance of core *Arabidopsis* plasmodesmal proteins in type I (four day old cultured cells) versus type II (seven day old cells) plasmodesmata. In *Arabidopsis* cultured cells, transition from type I to type II plasmodesmata is associated with a change in ER-PM contact site architecture, from very tight contact (~3 nm) with no visible cytoplasmic sleeve (type I) to larger ER-PM distance (lG nm to more) with an electron lucent cytosolic sleeve and sparse spoke-like elements (type II)(Nicolas *et al*, 20l7a). We analysed the plasmodesmata proteome from four days old cultured cells where type I plasmodesmata represent 7G% of the total plasmodesmata population and at seven days where this proportion is reversed and type II become predominant(Nicolas *et al*, 20l7a). Results show that 47 proteins from the plasmodesmata core proteome are differentially enriched at either type I or type II plasmodesmata, including all members of MCTPs (in bold), which are more abundant (l.4 to 3.6 folds) in type I plasmodesmata. Numbers in brackets correspondent to the protein numbering in Suppl. Table 1.

**Supplementary Figure 3.**
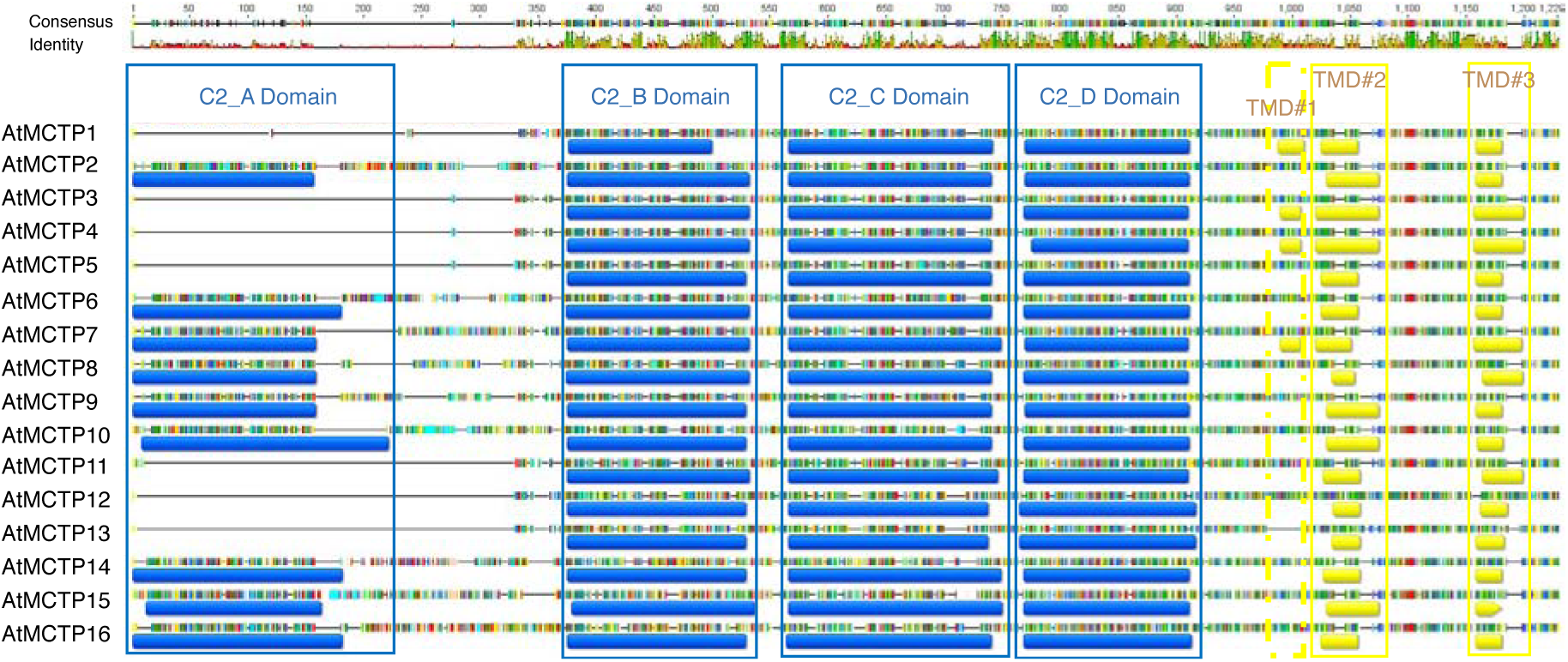
Domain organisation of the *Arabidopsis* MCTP protein family. Alignment of the 16 MCTP proteins of *A. thaliana*. C2 domains are represented in blue and transmembrane domains (TMD) in yellow. Each coloured vertical bars represents specific amino acid. The consensus sequence and the percentage of identity are represented on the top of the alignment. Note that for every MCTP member the C2 domains were individually delimitated using a combination of prediction methods (see M&M for details).

**Supplementary Figure 4.**
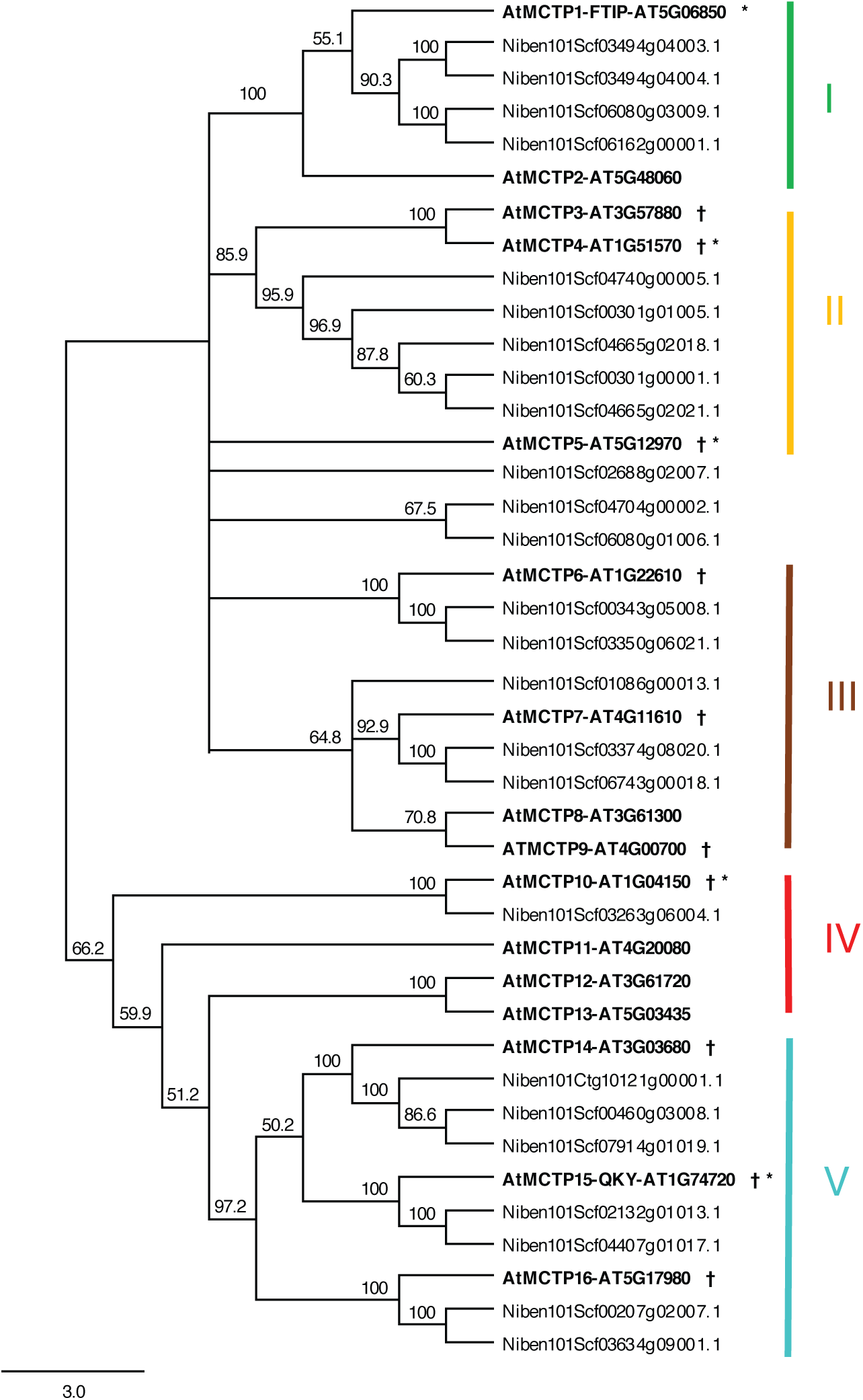
Phylogenetic tree of *A. thaliana* and *N. benthamiana* MCTP proteins. Amino acid sequences of MCTP family from *A. thaliana* and *N. benthamiana* were aligned with CLUSTALW(Thompson *et al*, 1997). The resulting alignment was adjusted manually and used to construct an unrooted phylogenetic tree using the neighbour-joining algorithm with Geneious 8.0.5 (https://www.geneious.com). Bootstrap values for 1000 re-samplings are shown on each branch.† indicates the MCTP members enriched in the plasmodesmata proteome and * indicates the MCTP members enriched in type I plasmodesmata. The five clades defined in Liu *et al*. 2017(Liu *et al*, 2017) are indicated from I to V.

**Supplementary Figure 5.**
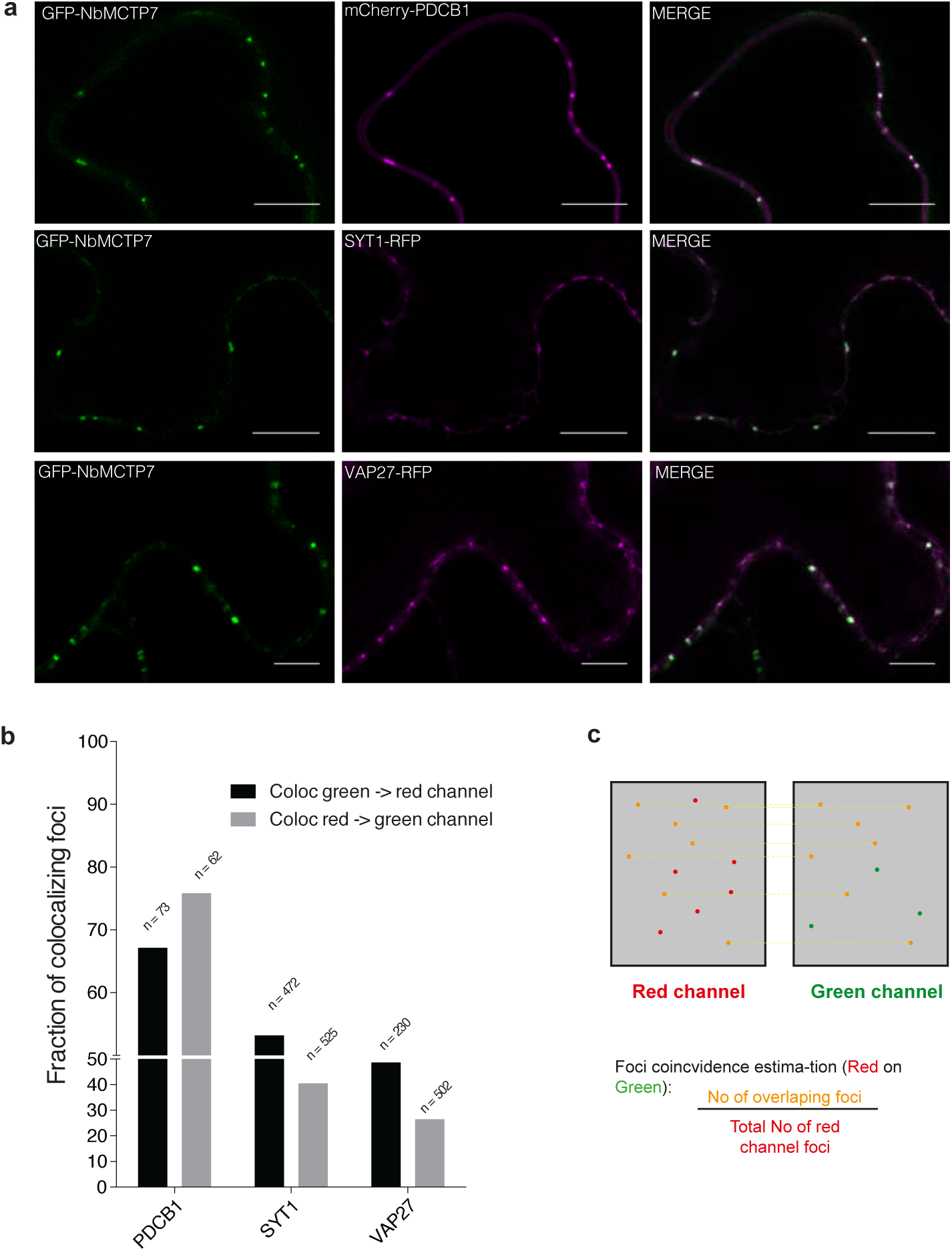
NbMCTP7 only partially co-localise with peripheral ER-PM contact sites. (a) Co-localisation between GFP-NbMCTP7 with mCHERRY-PDCB1 and two well-established markers of peripheral ER-PM contact sites, VAP27.1(Wang *et al*, 2016) and SYT1(Peréz-Sancho *et al*, 2015a; Levy *et al*, 2015), in *N. benthamiana* epidermal cells visualised by confocal microscopy. Scale bars, 10 μm. (b) Plot of the coincidence ratios. “Coloc green -> red channel” depicts the proportion of foci in the green channel overlapping with foci of the red channel over the total number of foci in the green channel. “Coloc red -> green channel” depicts this same proportion but of the red foci over the green foci. Coefficients range from 0 (complete exclusion) to 100% (complete colocalization of all foci). N indicated is the number of foci counted over 10 images of a given condition acquired over multiple co-expression/imaging sessions. (c) Cartoon schematic on how the Coincidence ratio is calculated.

**Supplementary Figure 6.**
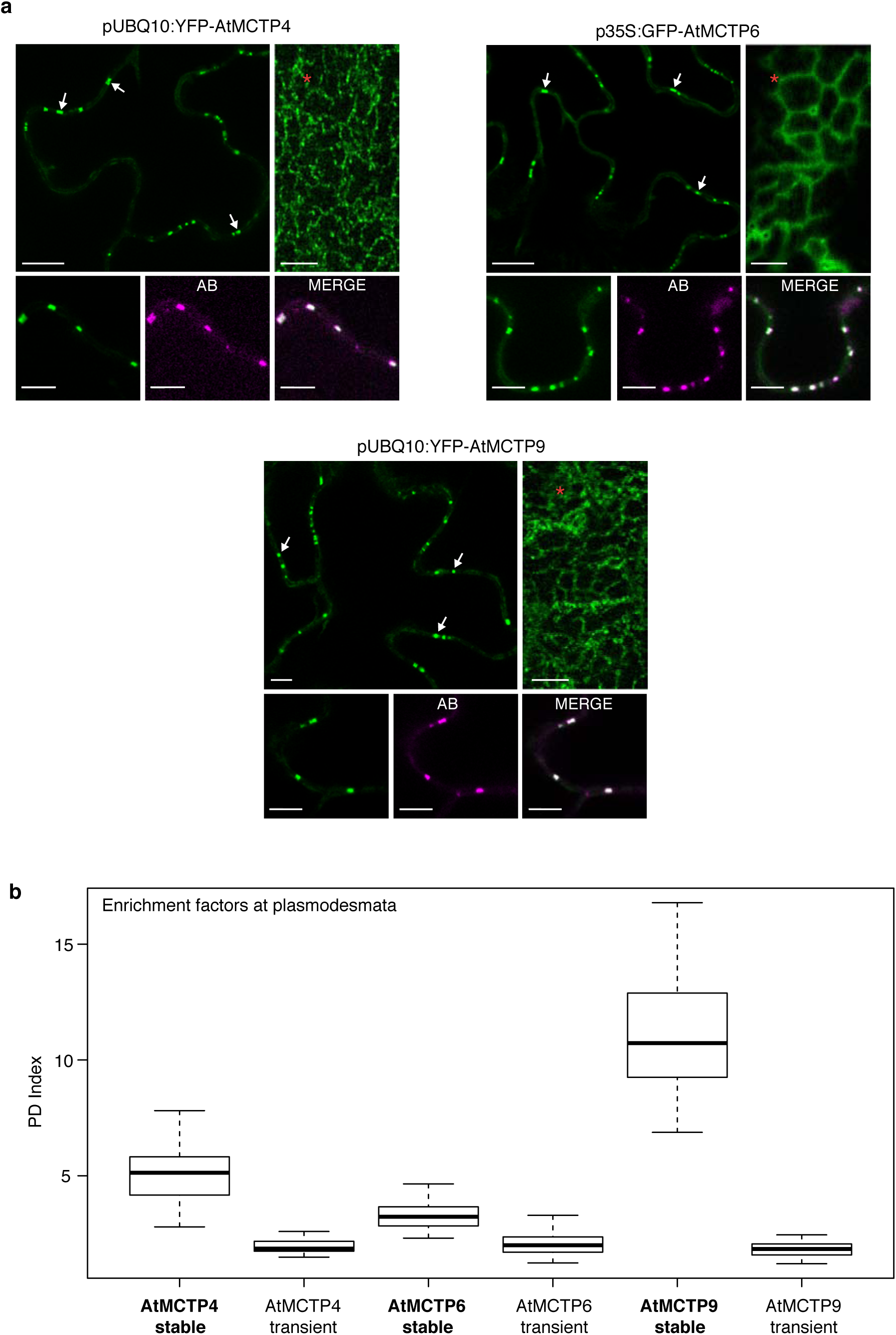
Subcellular localisation pattern of AtMCTP4, AtMCTP6 and AtMCTP9 when stably expressed in *Arabidopsis*. (a) Subcellular localisation of pUB10:YFP-AtMCTP4, 35S:GFP-MCTP6 and pUB10:YFP-AtMCTP9 in transgenic *Arabidopsis* epidermal cells showing typical plasmodesmata punctate pattern at the cell periphery (white arrows) and reticulated ER pattern at the cell surface (red stars). Plasmodesmal localisation was confirmed by aniline blue (AB) co-staining. Scale bars, 5 μm. (b) Plasmodesmata (PD) index of *Arabidopsis* MCTPs when either stably expressed transgenic Arabidopsis, or transiently expressed in *N. benthamiana*, showing consistently increased plasmodesmata association in transgenic lines.

**Supplementary Figure 7.**
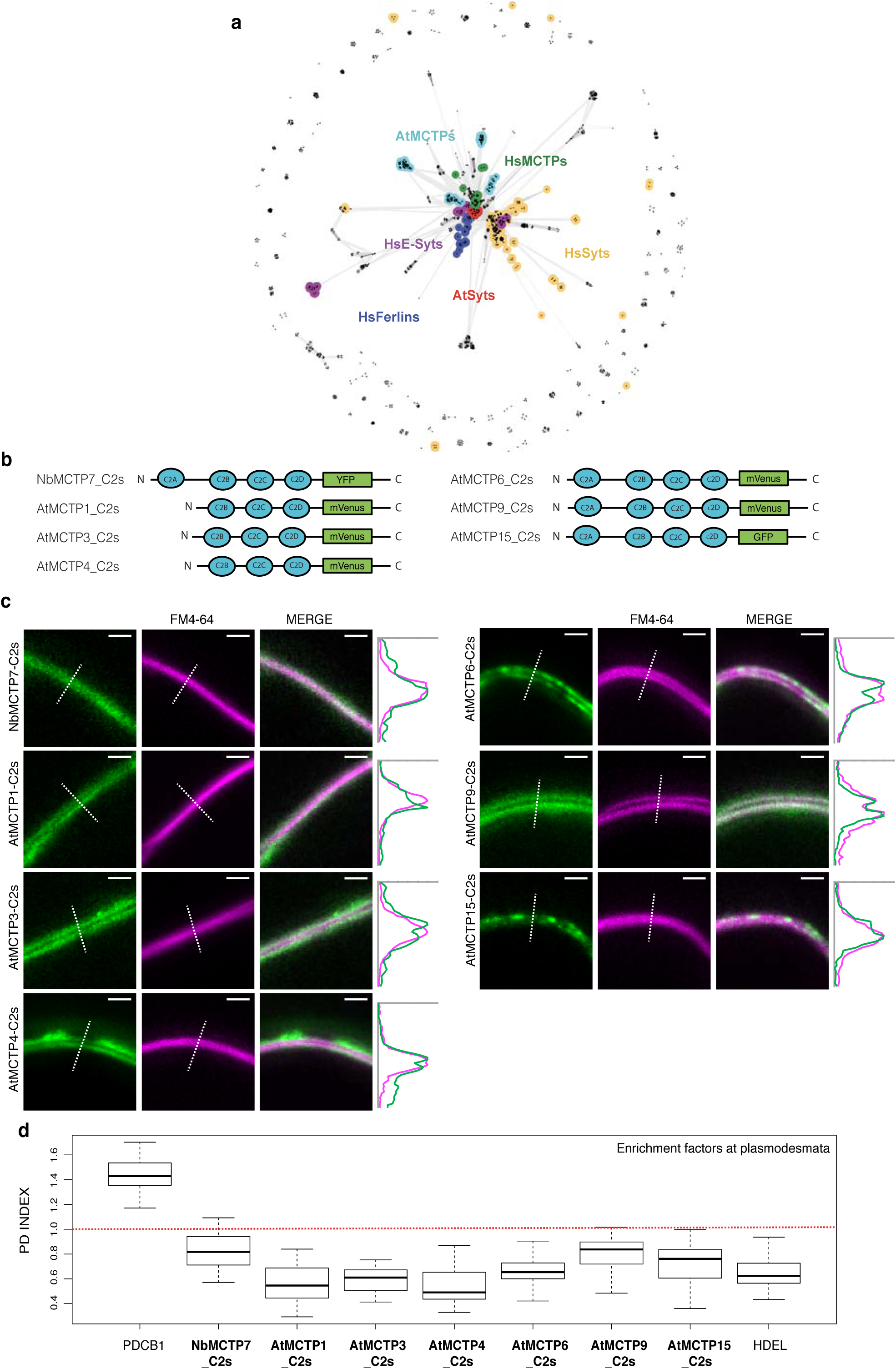
(a) Cluster map of human and *A. thaliana* C2 domains. Homologs of the four *A. thaliana* MCTP C2 domains were searched for in the human and *A. thaliana* proteomes using HHpred with a probability cut-off of 50% and with ‘No. of target sequences’ set to 10000. The obtained sequences were filtered to a maximum pairwise sequence identity of 100%, at a length coverage of 70%, using MMseqs2 (cite PMID: 29035372) to eliminate redundant sequences. The sequences in the filtered set, comprising almost all human and *A. thaliana C2* domains, were next clustered in CLANS based on their all-against-all pairwise sequence similarities as evaluated by BLAST P-values (PMID: 9254694). Clustering was done to equilibrium in 2D at a P-value cutoff of e-10 using default settings. In the map, dots represent sequences and line coloring reflects the strength of sequence similarity between them; the darker a line, the lower the P-value. Proteins not discussed in the manuscript are not colored. (b-d) The C2 blocks (C2A-D or C2B-D) of AtMCTP1, 3, 4, 6, 9, 15 and NbMCTP7 were tagged at their C-terminus with a fluorescent tag and expressed transiently in *N. benthamiana* leaves under moderate ubiquitin 10 promoter. b, Schematic representation of truncated MCTPs tagged with a fluorescent tag. c, Localisation of truncated AtMCTP1, 3, 4, 6, 9, 15 and NbMCTP7 C2 blocks (MCTP-C2s) in *N. benthamiana* epidermal cells by confocal microscopy. The PM was stained using short-term (up to 15 min) FM4-64 staining (magenta). Intensity plots are shown for each co-localisation pattern. When expressed in epidermal cells, MCTP-C2s-YFP constructs only partially associate with the PM and cytosolic localisation is also apparent. Scale bars, 5 μm. d, The PD index of individual truncated MCTP_C2s constructs is below 1 (red dashed line), indicating loss of plasmodesmata localisation.

**Supplementary Figure 8.**
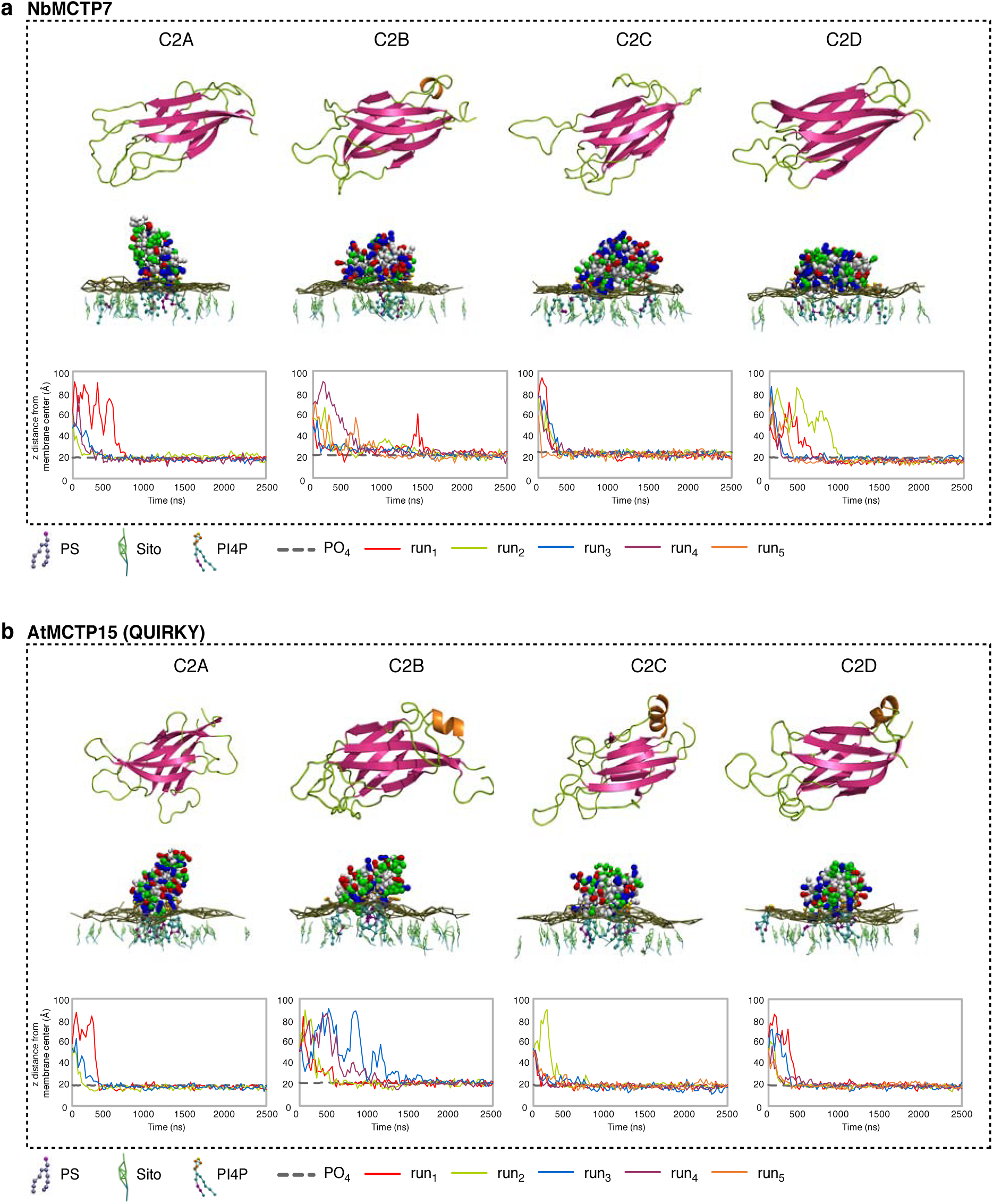
Membrane docking of NbMCTP7 and AtMCTP15/QKY C2 domains on a PM-like membrane. In (a) and (b); Top: 3D-atomistic model of the individual AtMCTP4 C2 domains. Beta strands are shown in pink, loops in green and alpha helices in orange. Bottom: molecular dynamics of individual NbMCTP7 (a) and AtMCTP15/QKY (b) C2 domains with phosphatidylcholine (PC), phosphatidylserine (PS), sitosterol (Sito) and phosphoinositol-4-phosphate (PI4P) (PC/PS/Sito/PI4P 57:19:20:4) biomimetic lipid bilayer. The plots show the minimal distance between the protein’s closest residue to the membrane and the membrane center, over time. The membrane’s phosphate plane is represented by a PO_4_ grey line on the graphs and a dark green meshwork on the simulation image captures (above graphs). For individual C2 domain, the simulations were repeated three to five times (runs 1-5). C2 membrane docking was only considered as positive when a minimum of three independent repetitions showed similarly stable interaction with the membrane. All C2 domains of NbMCTP7 and AtMCTP15/QKY show membrane interaction with a “PM-like” membrane composition, mainly due to the presence of PI4P. The amino acid colour code is as follow: red, negatively charged (acidic) residues; blue, positively charged (basic) residues; green, polar uncharged residues; and white, hydrophobic residues.

**Table S1.**
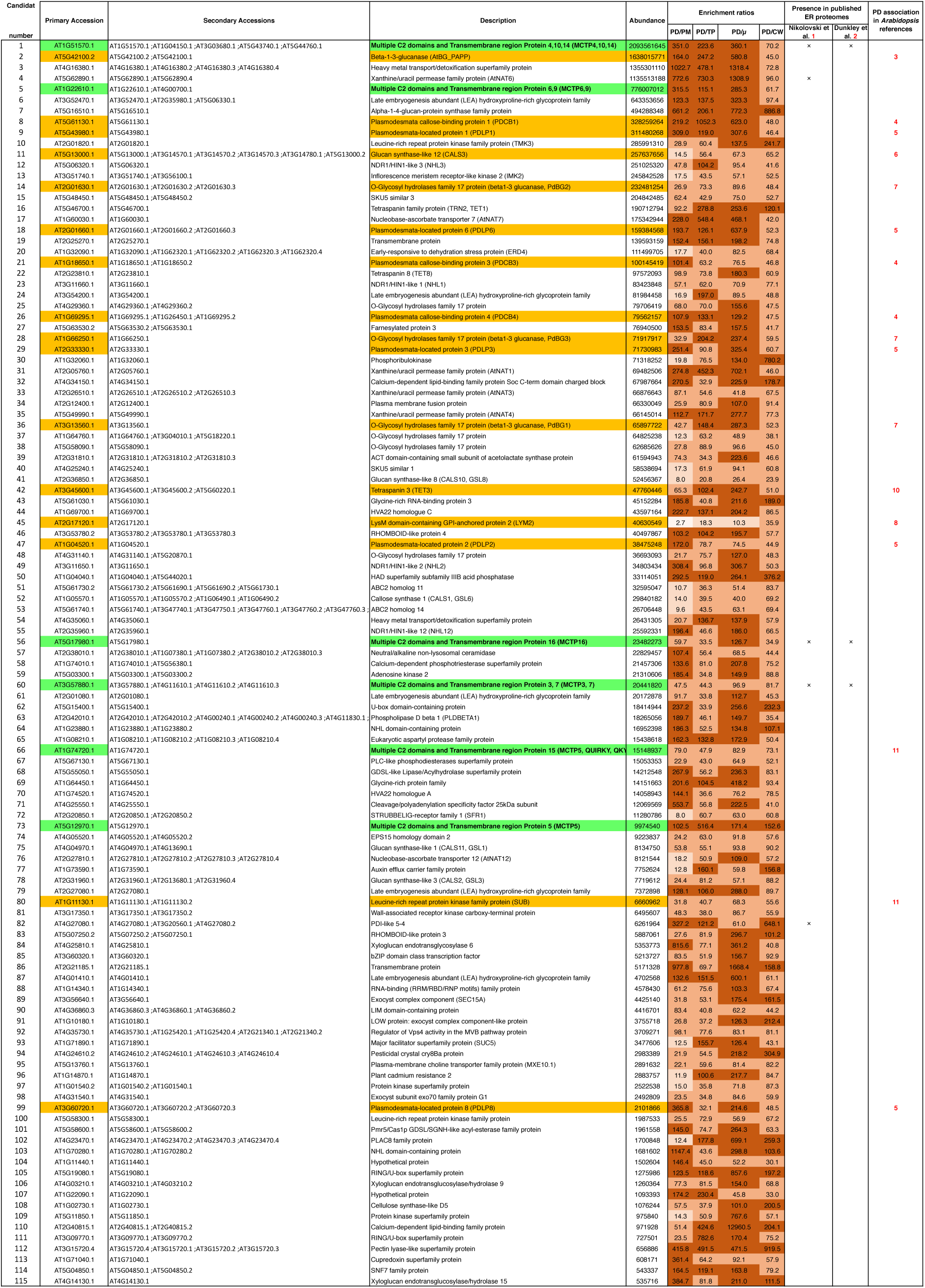
**Proteins of the core *Arabidopsis* plasmodesmata proteome** Label-free quantitation strategy was used to determine the relative abundance of proteins in the plasmodesmata (PD) fraction *versus* contaminant subcellular fractions namely, the PM, total extract (TP), microsomes (μ) and cell wall (CW), see Methods for details. Only proteins presenting minimum enrichment ratios of 8, 40, 30 and 30 in plasmodesmata *versus* PM, TP, microsomal and CW fractions, respectively were selected. Previously characterised plasmodesmal proteins are in orange and MCTP members in green. First row indicates the main accession and second row all possible isoforms potentially identified. The different shades (light to dark) of brown represent different enrichment levels (0-10; 10-20; 20-100 and above 100)

**Table S2.**
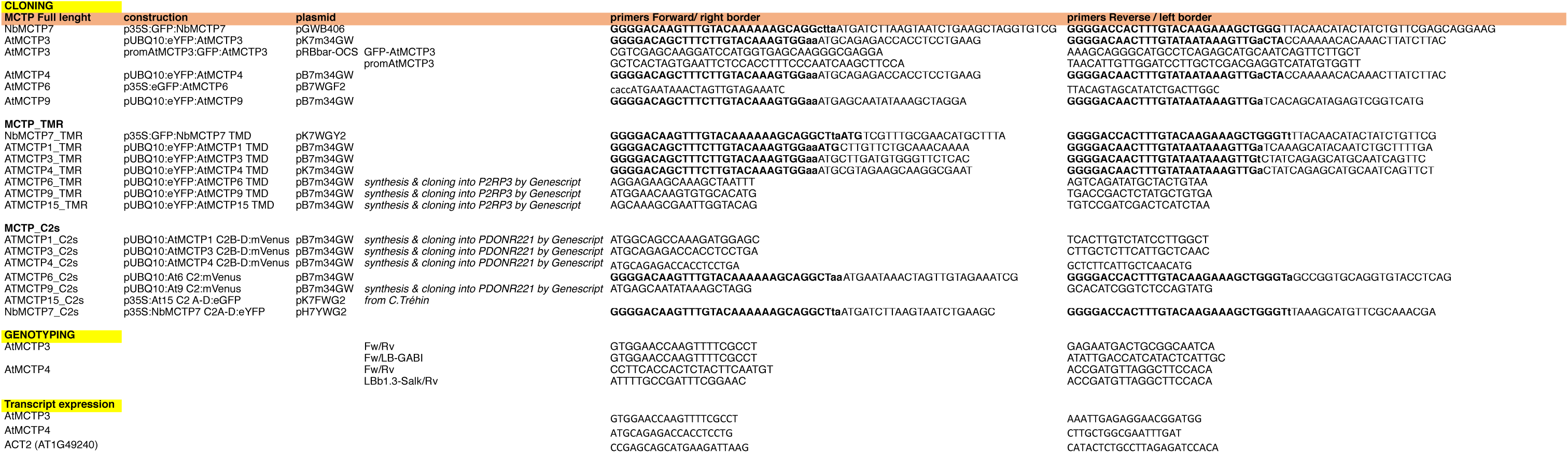
Primers used for MCTP cloning

**Movie S1.** Confocal time lapse imaging of 35S:GFP-NbMCTP7 in *N. benthamiana* epidermal leaves. One image every 0.2 seconds.

**Movie S2.** Confocal time lapse imaging of AtMCTP4:GFP-AtMCTP4 in transgenic *Arabidopsis* epidermal leaves. One image every 0.2 seconds.

**Movie S3.** Docking of the C2B, C2C and C2D domains of AtMCTP4 on a “PM-like” membrane (see Fig. 5), containing phosphatidylcholine (PC), phosphatidylserine (PS), sitosterol (Sito) and phosphoinositol-4-phosphate (PI4P) in the following ratio: PC/PS/Sito/PI4P 57:19:20:4. Please note that 0.5μs out of total (2.5μs) simulation is shown (moment of docking). The amino acid colour code is as follow: red, negatively charged (acidic) residues; blue, positively charged (basic) residues; green, polar uncharged residues; and white, hydrophobic residues. The lipid colour code is as follow: PC is depicted as light-pink polar heads and grey acyl chains, PS is depicted as dark-pink polar heads and light-purple acyl chains, PI4P is depicted as orange (inositol ring) and yellow (phosphate 4) polar heads and light-blue acyl chains and sitosterol is light-green.

